# Molecular Characterization of the Sea Lamprey Retina Illuminates the Evolutionary Origin of Retinal Cell Types

**DOI:** 10.1101/2023.12.10.571000

**Authors:** Junqiang Wang, Lin Zhang, Martina Cavallini, Ali Pahlevan, Junwei Sun, Ala Morshedian, Gordon L. Fain, Alapakkam P. Sampath, Yi-Rong Peng

## Abstract

The lamprey, a primitive jawless vertebrate whose ancestors diverged from all other vertebrates over 500 million years ago, offers a unique window into the primordial formation of the retina. Using single-cell RNA-sequencing, we characterized retinal cell types in lamprey and compared their molecular differentiation and regulatory networks with those in mouse and other jawed vertebrates. Our analysis revealed six cell classes and 74 distinct cell types. We discovered multiple conserved cell types shared between jawless and jawed lineages, including notably rods and cones, ON and OFF bipolar cells, and starburst amacrine cells. The conservation of these cell types indicates their emergence early in vertebrate evolution, highlighting the primal designs of retinal circuits for the rod pathway, ON-OFF discrimination, and direction selectivity. In contrast to this evidence for conservation, the pathways of diversification for amacrine cells and retinal ganglion cells appear to have distinctly diverged between the two lineages. Furthermore, we inferred master regulators in specifying retinal cell classes in both lamprey and macaque and identified common regulatory elements across species, underscoring the ancestral nature of the molecular origins governing retinal cell classes. Altogether, our characterization of the lamprey retina illuminates the evolutionary origin of visual processing in the retina.

## Introduction

The complex structure of the vertebrate nervous system arises from a distinct array of cell types present within each processing center. Much effort has recently been given to classifying cell types and describing their molecular differences^1,2^. Our understanding of neuronal cell-type specification is nowhere more advanced than in the vertebrate retina, where the description of its layered structure containing photoreceptors, interneurons, and ganglion cells dates back to the pioneering work of Ramón y Cajal^3^. More recent experimentation has identified over 100 cell types in mammals distributed into six cell classes, and much is known about the anatomy, development, physiology, and pharmacology of these different cell types ^4,5^. It is striking that the fundamental structure of the retina is conserved across all vertebrate lineages ^6,7^, which seems to indicate an ancient origin; but the cellular and molecular blueprint for the evolutionary formation of the retina remains largely unknown.

We hoped to shed some light on the evolution of the retina by studying the lamprey, a jawless vertebrate (agnathan) whose progenitors diverged from other vertebrates in the Cambrian over 500 million years ago (Fig. 1a)^8,9^. The eye of the adult lamprey shows remarkable similarities in structure to the conserved camera eye of jawed vertebrates, with a cornea, lens, pigmented epithelium, and retina with laminar structure^10,11^. Moreover, the lamprey retina shares key anatomical features with higher vertebrates, containing three cellular layers interconnected within two synaptic plexiform layers (Fig. 1b)^11,12^; and it is duplex, with functionally distinct rods and cones^13,14^. There are, however, several remarkable differences, which may suggest a primitive state for the lamprey retina. First, the three-layer retina is acquired only after the lamprey undergoes metamorphosis^12,15,16^. Second, the rods in the lamprey are morphologically similar to cones, with an outer segment consisting of invaginating lamellae continuous with the plasma membrane^10,17^. Last, retinal ganglion cells are located on both sides of the inner plexiform layer, with their axons forming the optic fiber layer between the inner plexiform layer and the inner nuclei layer (Fig. 1b), instead of at the vitread margin of the retina as in jawed vertebrates (Fig. 1b).

**Figure 1.**
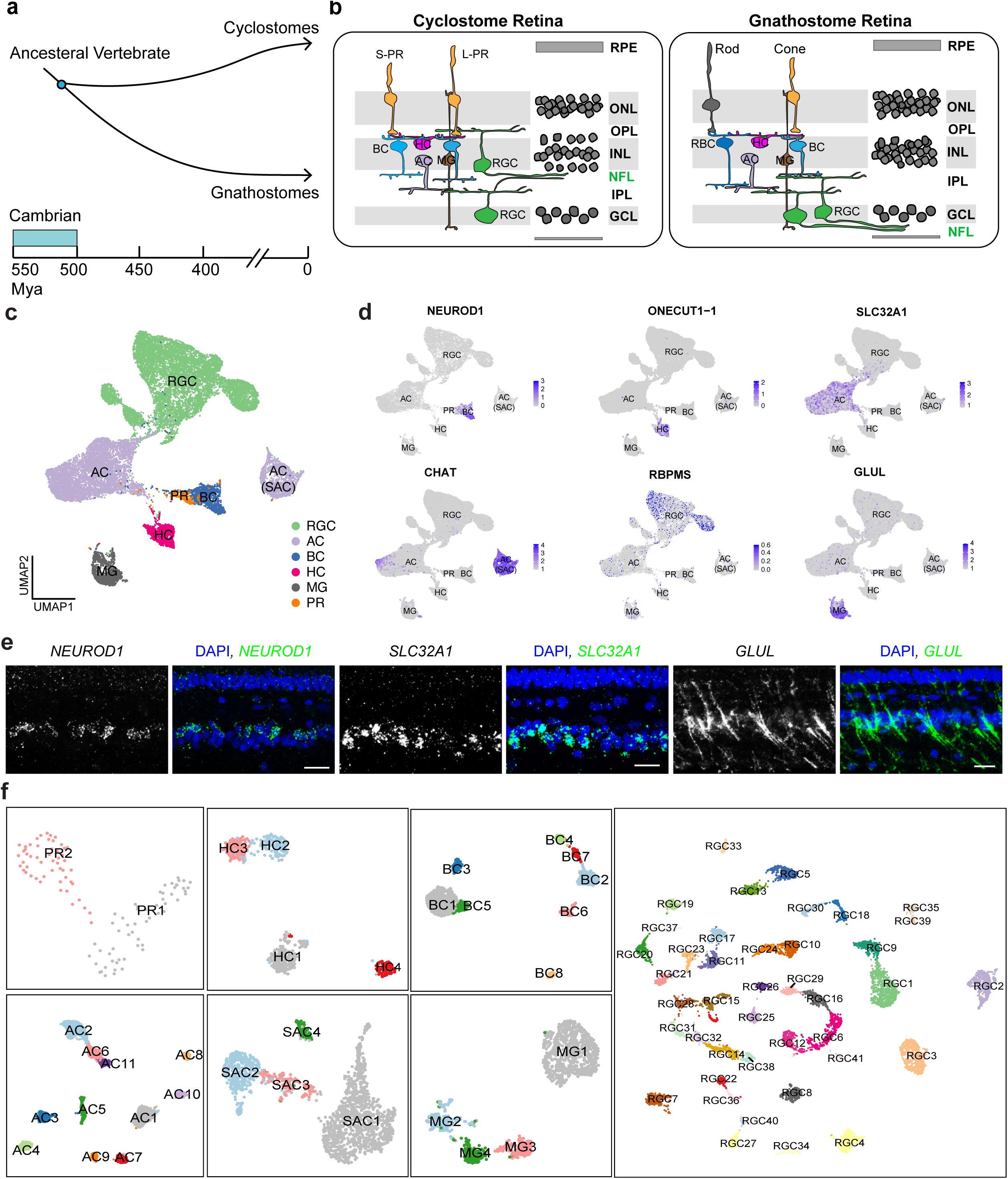
**Cell atlas of the adult lamprey retina** (a) Evolutionary history of vertebrates illustrating the split between cyclostomes (jawed vertebrates) and gnathostomes (jawless vertebrates) after vertebrate appearance during the Cambrian period over 500 million years ago (MYA). (b) Sketches of cellular arrangement and retinal circuitry in gnathostome (left) and cyclostome, particularly lamprey, (right). A major difference is the position of NFL (highlighted in green) between the two groups. (c) Uniform Manifold Approximation and Projection (UAMP) visualization of six lamprey cell classes, derived from a dataset of 21,474 cells. (d) Feature plots displaying the expression of canonical markers within individual cell classes. (e) Fluorescence in situ hybridization (FISH) validation of *NEUROD1* in bipolar cells, *SLC32A1* in amacrine cells, and *GLUL* in Müller glia. Scale bar, 20 μm. (f) UMAP visualizations of various cell types within each of cell classes, with SACs separated from the other ACs. Abbreviations: AC, amacrine cell; SAC, starburst amacrine cell; BC, bipolar cell; HC, horizontal cell; MG, Müller glia; PR, photoreceptor; RGC, retina ganglion cell; GCL, ganglion cell layer; IPL, inner plexiform layer; INL, inner nuclear layer; NFL, retinal nerve fiber layer; ONL, outer nuclear layer; OPL, outer plexiform layer.

High throughput single-cell RNA-sequencing (scRNA-seq) has become a pivotal tool for characterizing cell types in the nervous system from various vertebrate and invertebrate species^18–21^. This approach offers a comprehensive assessment of genomic activity within individual cell types. By comparing gene-expression patterns of cell types among species, we can construct homologous relationships and trace the evolutionary trajectory of cell-type differentiation^22,23^. Extending this approach to the lamprey could provide a much deeper understanding of the evolution of cell types in the retina, reaching further back into the history of vertebrates. The comparison of cell types across a large phylogenetic distance nevertheless presents ongoing challenges, resulting from incomplete genome and gene annotation of non-model organisms, distinct adaptations of model species to their environments, and the loss of homologous genes during species divergence^21,24,25^. Moreover, cell types are specified by transcriptional programs, which control genomic accessibility in each cell type^26^. Evaluating the conservation of these regulatory mechanisms between lamprey and jawed vertebrates can shed light on the genetic underpinnings of cell-type evolution in the vertebrate retina.

In this study, we used scRNA-seq to characterize the cell types in the adult retina of the sea lamprey (*Petromyzon marinus*). To facilitate cross-species comparison, we constructed a retina-specific transcriptome, which enhances the annotation of coding genes specific to lamprey retina tissue. Our scRNA-seq data revealed a total of 74 distinct cell types from all six cell classes. Comparative analyses of lamprey cell types against those from other vertebrate species revealed multiple conserved cell types, including, among others, rod and cone photoreceptors, rod bipolar cells, AII-like amacrine cells, Type1 and Type2 horizontal cells, ON and OFF starburst amacrine cells, and direction-selective ganglion cells. These results indicate that the foundational circuitry for specific features of light detection and signal integration in the retina emerged in the very earliest vertebrates, likely before the split of lamprey and other cyclostomes from the vertebrate lineage. To investigate shared genetic regulatory elements across the phylogenetic tree, we developed network-based methodologies to infer the activities of specific proteins, including transcription factors (TFs), transcription cofactors (coTFs), and surface signaling proteins. We found distinct levels of conservation among cell classes, grounded by their utilization of identified regulatory programs; and we have identified genetic programs that are likely inherent to the whole vertebrate lineage. Our work has provided new insight into the evolution of retinal cell types and the mechanisms by which type specifications were first established, which formed the basis of light detection in the vertebrate eye.

## Results

### Cell atlas of the adult sea lamprey retina

To facilitate cell and gene discovery, we used TruSeq to generate a retina-specific transcriptome of the lamprey (see Methods). We first compared public lamprey genome and annotation files from Ensembl (Pmarinus_7.0) and NCBI (kPetMar1.pri)^27^. Between the two, kPetMar1 performed better, because the mapping percentage of reads for genome, exons, and transcriptome was higher when we used it for alignment (Supplementary Fig. 1a). We thus built a retina-specific transcriptome based on the kPetMar1.pri version and named it as “NCBI+TruSeq.” Using NCBI+TruSeq transcriptome files, we significantly improved mapping metrics (Supplementary Fig. 1a). Notably, the newly updated transcriptome showed improved gene-body definitions for multiple genes that are crucial for the physiological function of retinal cells. For example, the red-opsin gene was updated with an additional 3’ exon, where abundant reads from both TruSeq and single-cell RNA-seq were aligned (Supplementary Fig. 1b). Although the NCBI assembly kPetMar1.pri is a chromosome-level genome assembly, over 60% of annotated genes were named with a “LOC” number without a gene symbol. We further annotated these LOC genes to correlated gene symbols based on the information of their gene products (See Methods). The final reference transcriptome has nearly 80% of its genes with specific gene symbols, thus greatly improving our ability to classify cell types and characterize molecular profiles of lamprey retina cells (Supplementary Fig. 1c, Supplementary Table 1).

After mapping scRNA-seq reads to the retina-specific transcriptome, we obtained 21,474 high-quality transcriptomes from individual cells. Using an unsupervised clustering method, we identified six main cell classes: photoreceptors (PR), horizontal cells (HC), bipolar cells (BC), amacrine cells (AC), retinal ganglion cells (RGC), and Müller glia (MG) (Fig. 1c)^28^. No bias was observed associated with biological replicates (Supplementary Fig. 2a, 2b). Each lamprey cell class showed the expression of a similar set of marker genes, which are also expressed in the mouse or macaque retina (Fig. 1d)^29,30^. The expression of these canonical cell-class markers was confirmed by fluorescence *in situ* hybridization in the lamprey retina (Fig. 1e). We further clustered cell types from individual cell classes, with ACs separating into starburst amacrine cells (SACs) and other AC subclasses (Fig. 1f). We identified a total of 2 PR, 4 HC, 8 BC, 4 SAC, 11 non-SAC AC, 41 RGC, and 4 MG clusters (Fig. 1f). These cell types were equivalently identified from either replicate (Supplementary Fig. 2c). Using the hierarchical clustering^30^, we found that each cell type from the same cell class is more closely correlated from within their class than from outside of their class (Supplementary Fig. 2d). Thus, the lamprey retina is composed of at least 74 cell types with clear molecular distinction.

### Conserved retinal cell types between the lamprey and jawed species

The number of diverse cell types identified in the lamprey is comparable to that seen in higher vertebrate species^4,5^, suggesting that cell-type diversification might have originated with the very earliest of vertebrates. To explore this hypothesis in greater detail, we attempted to compare lamprey cell types to those in jawed vertebrate species. We employed two ways to achieve this comparison. 1) We integrated lamprey cell types in individual classes with cell types in selected species. 2) We used transcriptomic mapping based on multiclass classification frameworks to identify the closest related cell types in jawed species, relying primarily on mouse and chicken since their retinal-cell atlases have been well defined and validated. We found examples of strikingly conserved cell types between the lamprey and these two species, as well as interesting differences.

#### Photoreceptors

We identified two lamprey photoreceptors (PR): PR1 and PR2 (Fig. 2a). The expression of rhodopsin in PR1 and red-opsin in PR2 identify these morphologically defined “short” and “long” PRs as a rod and a single type of cone (Fig. 2b, 2c and Supplementary Fig. 3), in agreement with results from physiological recordings and molecular validations^13,14,31–33^. The functional differences between rods and cones are achieved through the use of distinct molecules in their phototransduction cascades^34,35^. Interestingly, we found strikingly similar differences in the phototransduction cascades between lamprey PR1 and PR2 (Fig. 2b)^11^. We further confirmed that the G-protein subunit alpha transducin 1 (GNAT1) is expressed in PR1, while GNAT2 is expressed in PR2 (Fig. 2C)^36^.

**Figure 2.**
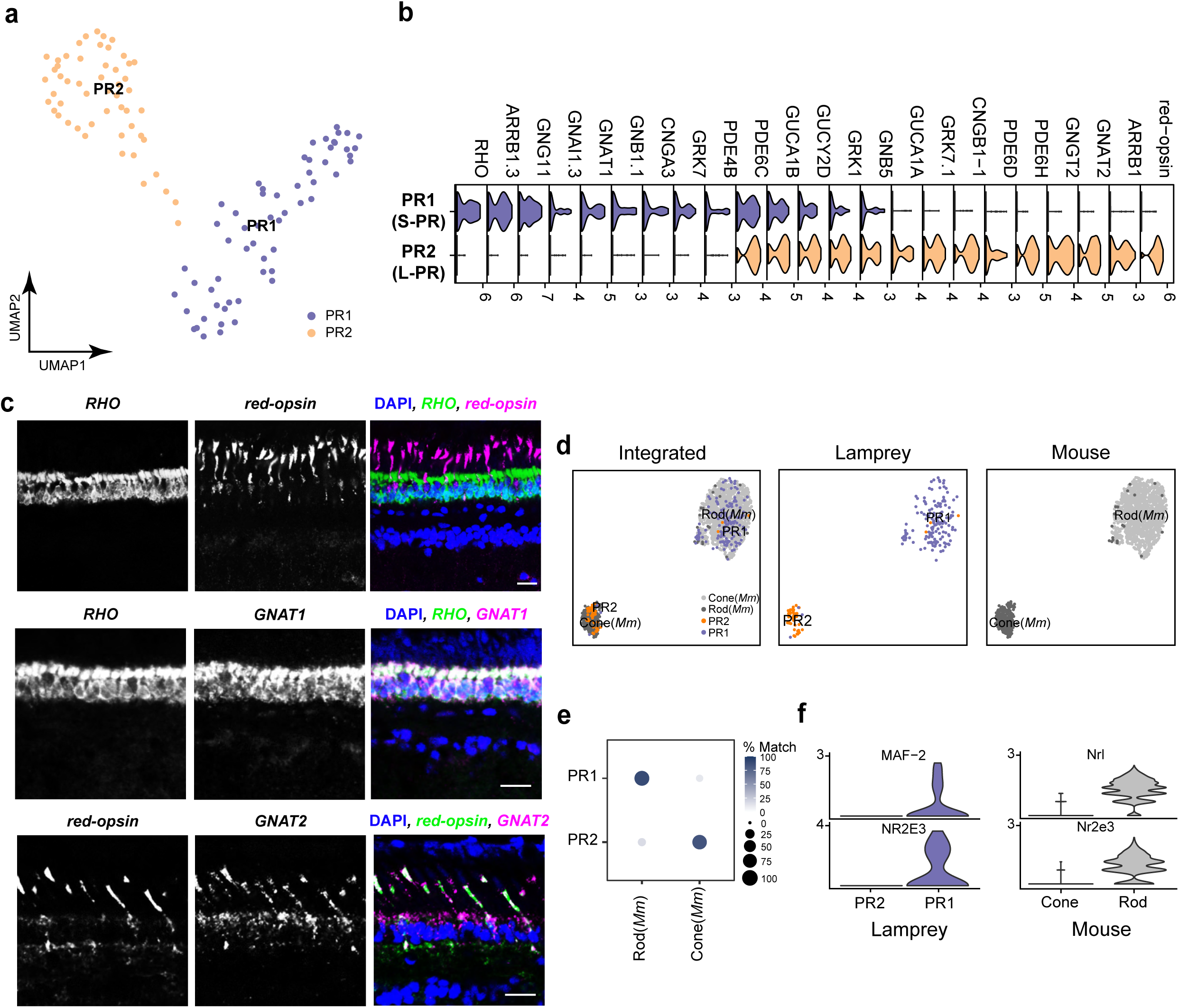
**Classification of lamprey PR types and their comparison with mouse PR types** (a) UMAP visualization of two lamprey PR types. (b) Stack violin plot showing distinct gene-expression patterns in the phototransduction cascade between PR1(S-PR) and PR2(L-PR). S-PR, short-photoreceptor; L-PR, long-photoreceptor. (c) FISH validations showing an exclusive expression of *Rhodopsin (RHO)* and *red-opsin* between PR1 and PR2 (upper panel), co-expression of *RHO* and *GNAT1* in PR1 (middle panel), and co-expression of *red-opsin* and *GNAT2* in PR2 (bottom panel). Scale bar, 20 μm. (d) Integration of lamprey and mouse PRs visualized with UMAP, with both integrated and species-specific clusters presented in separated UMAP plots. (e) Confusion matrix demonstrating the transcriptomic correspondence of PR types between lamprey and mouse (*Mus musculus,* Mm). Mouse PRs were used as the training dataset, while the lamprey PRs were used as the testing dataset. Circles and color gradients convey the percentage of cells from a mouse PR type assigned to a corresponding lamprey PR cluster. (f) Violin plots showing the expression of conserved transcription factors enriched in both lamprey PR1 and mouse rods.

To assess the molecular similarities between lamprey and mouse photoreceptors, we compared their transcriptomes. We integrated mouse PRs^37^ with lamprey PRs and identified two clusters through an unsupervised method (Fig. 2d). As expected, lamprey PR1 aligned well with mouse rods, while PR2 aligned with mouse cones (Fig. 2d). These findings are supported by transcriptomic mapping, which shows that PR1 corresponds to mouse rods and PR2 to mouse cones (Fig. 2e).

Different transcription factors distinguish rod from cone fate in the mouse retina, such as Nrl and Nr2e3 (Fig. 2f)^38–41^. Notably, we found that either the same gene (*NR2E3*) or a gene from the same NRL-MAF-family (*MAF-2*) is differently expressed in PR1 and PR2^42^. These findings, taken together, show that PR1 and PR2 are molecularly similar to rods and cones, and that their differentiation is likely regulated by at least partially conserved transcriptional mechanisms.

#### Bipolar cells

We identified 8 BC types (Fig. 3a). BCs are typically categorized into ON and OFF subclasses based on the expression of metabotropic and ionotropic glutamate receptors ^43^. By studying the genes encoding these channels, we found that lamprey BCs can be similarly classified into ON types expressing the group III metabotropic glutamate receptor GRM8.2, and OFF types expressing the kainate receptor GRIK2-2 or GRIK2 (Fig. 3b, Supplementary Fig. 4a). Notably, ON BC types also expressed *TRPM3.1*—metabotropic glutamate receptors often pair with transient receptor-potential cation channels (Fig. 3b, Supplementary Fig. 4a). Thus, lamprey BC types can be molecularly classified into ON and OFF subclasses. Unlike mammalian BCs, lamprey ON BCs expressed *SLC17A6*, a glutamate transporter in mammals specific to retinal ganglion cells (Fig. 3b, 3c)^44^.

**Figure 3.**
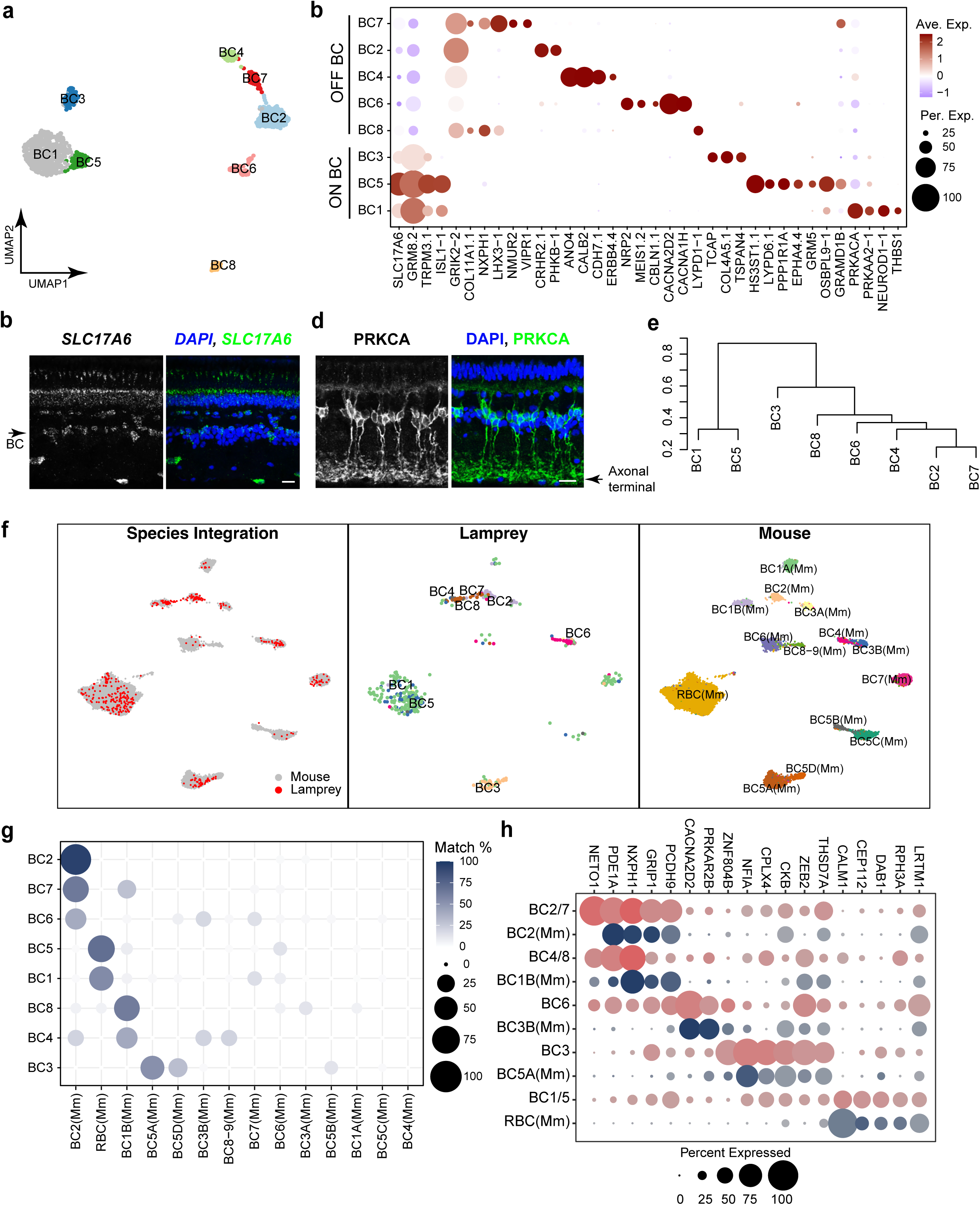
**Classification of lamprey BC types and their comparison with mouse BC types** (a) UMAP visualization of eight BC clusters in the lamprey retina. (b) Dot plot showing expression patterns of markers specific to ON and OFF BC subgroups, as well as markers unique to individual BC types. Abbreviations: Per. Exp, expression percentage; Ave. Exp, average expression. (c) FISH validation of SLC17A6 expression in multiple cell types in the lamprey, including BC (arrow). Scale bar, 20 μm. (d) Immunostaining with PRKCA in the lamprey retina showing the morphology of rod bipolar cells (RBCs), with axonal terminals of RBCs indicated by arrow. Scale bar, 20 μm. (e) Complete linkage agglomerative hierarchical clustering of BC types from correlation distance and complete linkage. The scale on the left indicates correlation distance. (f) Integration of lamprey and mouse BCs visualized with UMAP, with both integrated and species-specific clusters presented in separated UMAP plots. (g) Confusion matrix demonstrating the transcriptomic correspondence of BC types between lamprey and mouse. Mouse (Mm) BCs were used as the training dataset, while lamprey BCs were used as the testing dataset. (h) Dot plot showing expression patterns of conserved markers across corresponding BC types in lamprey and mouse (Mm).

Given that rods and cones are molecularly distinct in the lamprey retina, we explored the possibility of classifying BC types into rod BCs (RBCs) and cone BC types (CBCs)^45,46^. In mammals, protein kinase C alpha (PrkCa) and Gramd1b are markers for RBCs^30,45^. We found that *PRKACA* (an alias of *PRKCA*)^47^ is highly expressed in BC1, and *GRAMD1B* is highly expressed in BC5 and BC7. From the glutamate receptors expressed by these cells (Fig. 3b), we classified BC1 and BC5 as ON and BC7 as OFF. From hierarchical clustering and correlation-expression analysis, we determined that BC1 and BC5 are closely related and stand apart from the other types (Fig. 3e, Supplementary Fig. 4b). We suspect that both of these cells are ON rod BCs and may be related to the two types of ON rod BCs recently characterized in zebrafish retina^48^. Unlike mammals, lamprey have a rod bipolar cell that is hyperpolarizing, which could be BC7^49^. Using anti-PRKCA antibody to stain the lamprey retina, we found that PrkCa-positive BCs resemble mouse RBCs morphologically. The cell bodies of these BCs press against the outer plexiform layer while their axons innervate the innermost part of the inner plexiform layer (Fig. 3d). It is likely that these are the BC1 cells.

We next compared the transcriptomic profiles of lamprey BC types with those in mice to investigate which lamprey BC types are closest to mouse RBCs. We integrated mouse BCs with lamprey BCs. Interestingly, lamprey BC1 and BC 5 align well with mouse RBCs, while all lamprey OFF BC types align with mouse BC1B or BC2 types. Lamprey BC3 aligns with mouse BC5A and BC5D (Fig. 3f). We further used transcriptomic mapping to correlate lamprey BC types to the 15 mouse BC types. BC1 and BC 5 were consistently analogous to the mouse RBC, with the remainder of lamprey bipolar cells corresponding either to ON CBCs (BC5A or BC5D) or to OFF CBCs (BC1B or BC2) (Fig. 3g). We detected conserved markers shared between correlated mouse and lamprey BC types (Fig. 3h). Thus, our results underscore that, akin to mammals, lamprey BC types are primarily divided into RBCs (which in lamprey can be either ON or OFF), and ON-CBC and OFF-CBC groups. It is particularly noteworthy that lamprey RBCs might comprise as many as three types^49^.

#### Horizontal cells

We identified four HC types in the lamprey retina (Fig. 4a). Nearly all sea lamprey HCs express the melanopsin-like gene–*OPN4l.1* (Fig. 4b, 4c), similar to the expression pattern in river lampreys^50^. Like chicken and macaque Type I and Type II HCs, lamprey HCs can be divided into two subgroups, each exclusively expressing either *ISL1* or *LHX1* (Fig. 4b)^19,30^. We further integrated data from lamprey HCs with chicken HCs^19^. In the chicken retina, there are five HC types: HC1 and 3 are classified as Type I HCs, while HC 2/4/5 belong to Type II HCs. Our integrated data separate into two clusters: lamprey H1 and H4 align with chicken HC2/4/5, and lamprey H2 and H3 align with chicken H1/3 (Fig. 4d). Using the transcriptomic mapping method, we obtained a corresponding result mirroring the integrated pattern (Fig. 4e). Moreover, we identified conserved marker genes between lamprey HC 1/4 and chicken HC 2/4/5, as well as between lamprey HC 2/3 and chicken HC 1/3 (Fig. 4f). Therefore, the four HC types in the lamprey correspond to the Type I and Type II HCs found in chicken.

**Figure 4.**
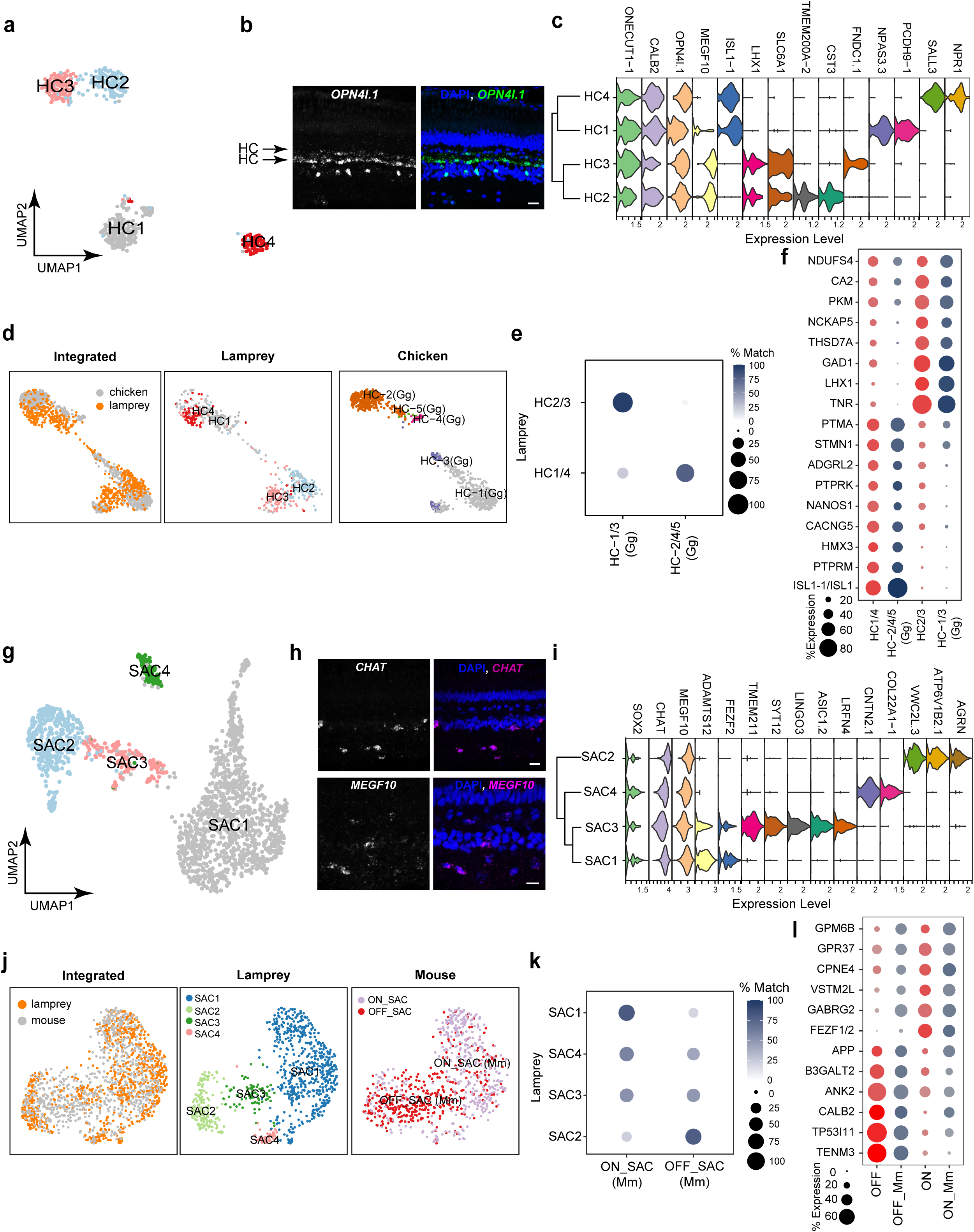
**Classification of lamprey HC and SAC types and their comparison with chicken or mouse types** (a) UMAP visualization of four lamprey HC types. (b) FISH validation of the expression of the melanopsin-like gene, *OPN4l.1*, in the lamprey retina. Arrows point to HC layers. (c) Stack violin plot showing expression patterns of common markers for all HC types and markers specific to individual HC types. The dendrogram on the left shows agglomerative hierarchical clustering of HC types. (d) Integration of lamprey and chicken HCs visualized with UMAP, with both integrated and species-specific clusters presented in separated UMAP plots. (e) Confusion matrix demonstrating transcriptomic correspondence of HC types between lamprey and chicken. Chicken (*Gallus gallus,* Gg) HCs were used as the training dataset, while lamprey HCs were used as the testing dataset. (f) Dot plot showing expression patterns of conserved markers across corresponding HC types in lamprey and chicken (Gg). (g) UMAP visualization of four lamprey SAC types. (h) FISH validation of the expression of *CHAT* and *MEGF10* in the lamprey retina. (i) Stack violin plot showing markers common to all SAC types and markers specific to individual SAC types. The dendrogram on the left gives agglomerative hierarchical clustering of SAC types. (j) Integration of lamprey and mouse SACs visualized with UMAP, with both integrated and species-specific clusters presented in separated UMAP plots. (k) Confusion matrix demonstrating transcriptomic correspondence of SAC types between lamprey and mouse. Mouse (Mm) SACs were used as the training dataset, while lamprey SACs were used as the testing dataset. (l) Dot plot showing the expression patterns of conserved markers across corresponding SAC types in lamprey and mouse (Mm).

#### Starburst amacrine cells

Starburst amacrine cells (SACs) are cholinergic cells in the retina, which are essential components of retinal circuits detecting the direction of motion^51–53^. SACs constitute ∼22% of all the amacrine cells in the lamprey retina, a proportion that is much greater than the 5% in mouse retina^54^. We identified four SAC clusters (Fig. 4g), all of which express canonical SAC markers, such as *SOX2*, *CHAT*, and *MEGF10* (Fig. 4h, 4i)^55,56^. Interestingly, SAC1 and SAC3 were found to express *FEZF2* (Fig. 4i), a gene from the same family as *Fezf1*, which in the mouse retina determines the fate and somatic position of ON SACs^57^. These findings suggest that SAC types in lamprey might also be classified into ON and OFF subgroups. In fact, when integrating data from lamprey SACs with mouse ones, we observed that lamprey SAC1 and SAC 4 aligned more closely with mouse ON SACs, while SAC2 and SAC3 are more closely aligned with mouse OFF SACs. Through transcriptomic mapping, we determined that among all the SAC types, SAC1 and SAC2 most closely correspond to mouse ON-SACs and OFF SACs (Fig. 4k). Moreover, they shared multiple conserved markers with mouse ON or OFF SACs (Fig. 4l). Therefore, our results indicate that lamprey SACs include both ON and OFF types, and that their fates may be determined at least in part by members of the *FEZ* gene family.

### The evolutionary diversification of AC and RGC types

The remarkable conservation across PR, BC, HC, and SAC types in the lamprey hints at a foundational program that is at the core of the structure of all vertebrate retinas. Our exploration now broadens to include the two other cell classes, ACs and RGCs, which across jawed vertebrates display the most heterogeneity in cell types. We found that the lamprey AC and RGC classes also contain the most heterogenous cell types compared to other classes. We related AC and RGC types from the lamprey to those of the mouse to understand the diversification paths of these two classes between jawless and jawed lineages.

#### Amacrine cells

In addition to the four SAC types, we identified 11 non-SAC AC types in the lamprey retina (Fig. 1f, Supplementary Fig. 5a). All AC types expressed a GABA transporter VGT3 (*SLC6A11*) as well as the glycine transporters GlyT1 (*SLC6A5*) or GlyT2 (*SLC6A9*) (Fig. 5a). This finding differs from the separate expression of these transporters in distinct GABAergic and glycinergic subgroups seen in jawed species^58^. Additionally, the number of AC types in the lamprey retina was less than one-quarter of AC types in chicken or mouse retina^19,58^, indicating that lamprey ACs are less diversified compared to jawed species. When we compared lamprey ACs to mouse or chicken AC types, either through integration or by using transcriptomic mapping methods, we observed a consistent pattern: multiple mouse or chicken AC types corresponded to a single lamprey type with over 50% matching percentage (Fig. 5b, Supplementary Fig. 5a-c). This correspondence suggests that AC types share similar origins between jawless and jawed lineages, but that these types may have further diversified into multiple sister types in jawed species. Notably, the AII-AC (AC3) in the mouse retina corresponds to lamprey AC8 with the highest confidence, sharing common markers including *CAR2* and *KCNMA1* (Fig. 5c). However, the gap-junction gene *GJD2* generally has low expression in lamprey ACs, and it is not expressed by AC8 (Fig. 5a). These findings suggest that an AII-like AC might have already been present in jawless species, but that these cells might not have had the same function or connectivity. The AC11 ACs in lamprey didn’t have good matching types in either chicken or mouse, suggesting that this type of cell may have evolved separately in lamprey or disappeared during the evolution of jawed vertebrates.

**Figure 5.**
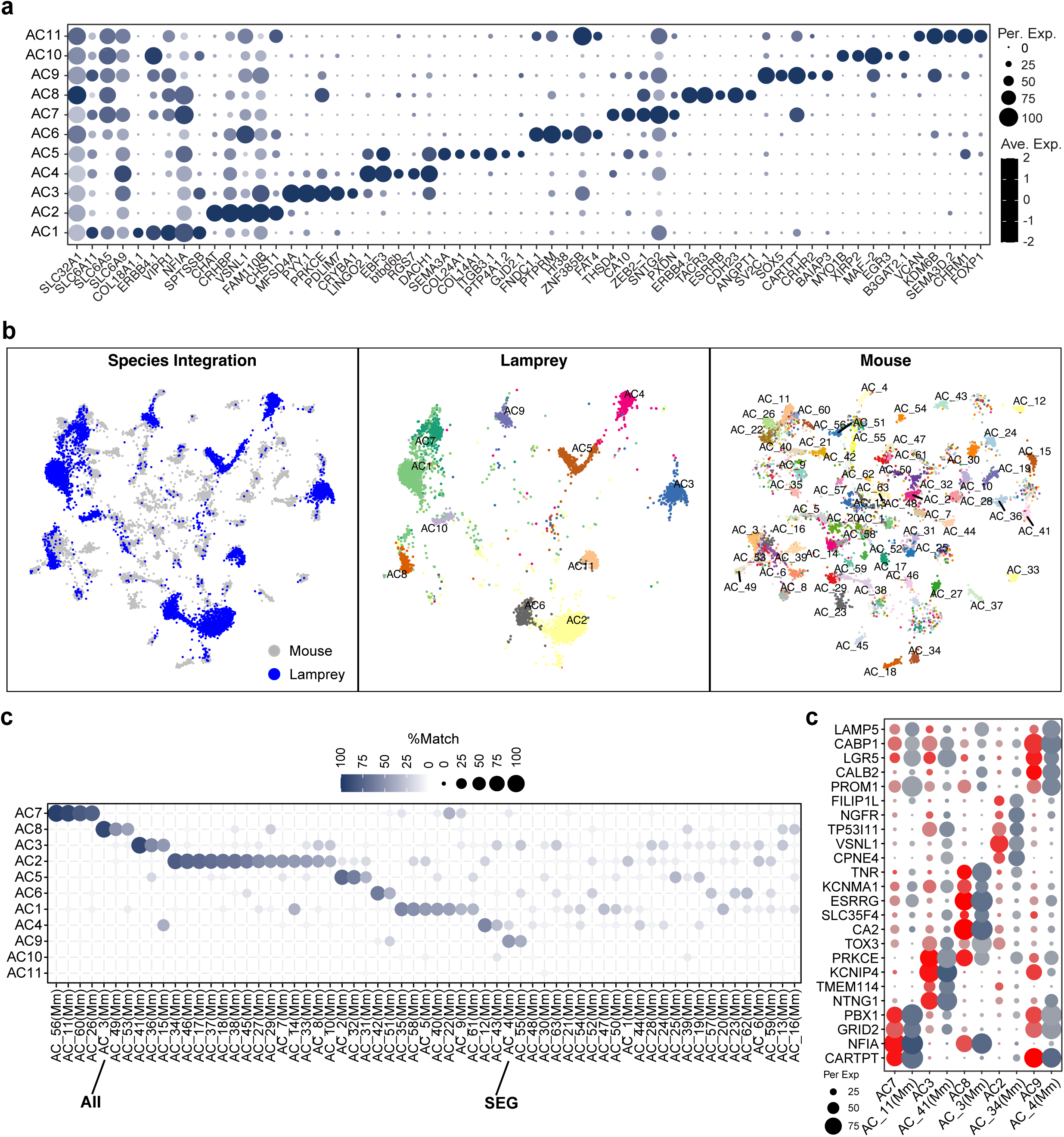
**Classification of lamprey AC types and their comparison with mouse AC types** (a) Dot plot showing the expression of genes for GABA transporters (*SLC32A1* and *SL6A11*) and glycine transporters (*SLC6A5* and *SLC6A9*) and marker genes for individual AC clusters. (b) Integration of lamprey and mouse ACs visualized with UMAP, with both integrated and species-specific clusters presented in separated UMAP plots. (c) Confusion matrix demonstrating transcriptomic correspondence of AC types between lamprey and mouse. Lamprey ACs were used as the training dataset, while mouse ACs were used as the testing dataset. Mouse AII AC and SEG AC are indicated based on Ref 58. (d) Dot plot showing expression patterns of conserved markers across corresponding AC types between lamprey and mouse.

#### Retinal ganglion cells

We identified 41 RGC types in the lamprey, a number similar to what has been reported in the chicken or mouse retina (Supplementary Fig. 6)^44,58,59^. Based on hierarchical clustering and the expression of shared marker genes, these 41 RGC types could be divided into seven subgroups: 1) *SEMA3A.1* and *LAMB2* positive RGCs, 2) *FOXP1* and *PRDM13* positive RGCs, 3) TMEM121-positive RGCs, 4) *PIEZO2.1* and PENK positive RGCs, 5) *VAT1* and *TAFA2.1* positive RGCs, 6) *OSBPL5* and *PDE11A* positive RGCs, 7) *NOS1* and *NR4A2* positive RGCs (Supplementary Fig. 6). We also identified two ipRGC types with expression of the *OPN4* paralog *OPN4l.1*^50^.

Morphological studies of lamprey RGCs identified two RGC subclasses, with 40% of RGCs located at the GCL layer, and the remaining 60% located at the inner nuclear layer^60,61^. The distinct somatic position and axonal efferent routes of RGCs in lamprey suggest significant divergence between jawless and jawed RGCs (Fig. 1b). Indeed, unlike the cross-species comparison of ACs, only a handful of RGC types matched well with mouse RGC types (Fig. 6a). Among the best matching types, we found that there are lamprey ganglion-cell types that correspond to OFF transient alpha RGCs (αOFF-T), ON-OFF direction-selective ganglion cells (ooDSGCs), J-RGCs, W3B RGCs, OFF sustain alpha RGCs (αOFF-S), and F-miniON-RGCs in mouse (Fig. 6b, 6c)^62–67^. It seems that many of these conserved RGC types may function to encode retinal motion that direct eye movements, considering the known functions of their counterparts in the mouse retina.

**Figure 6.**
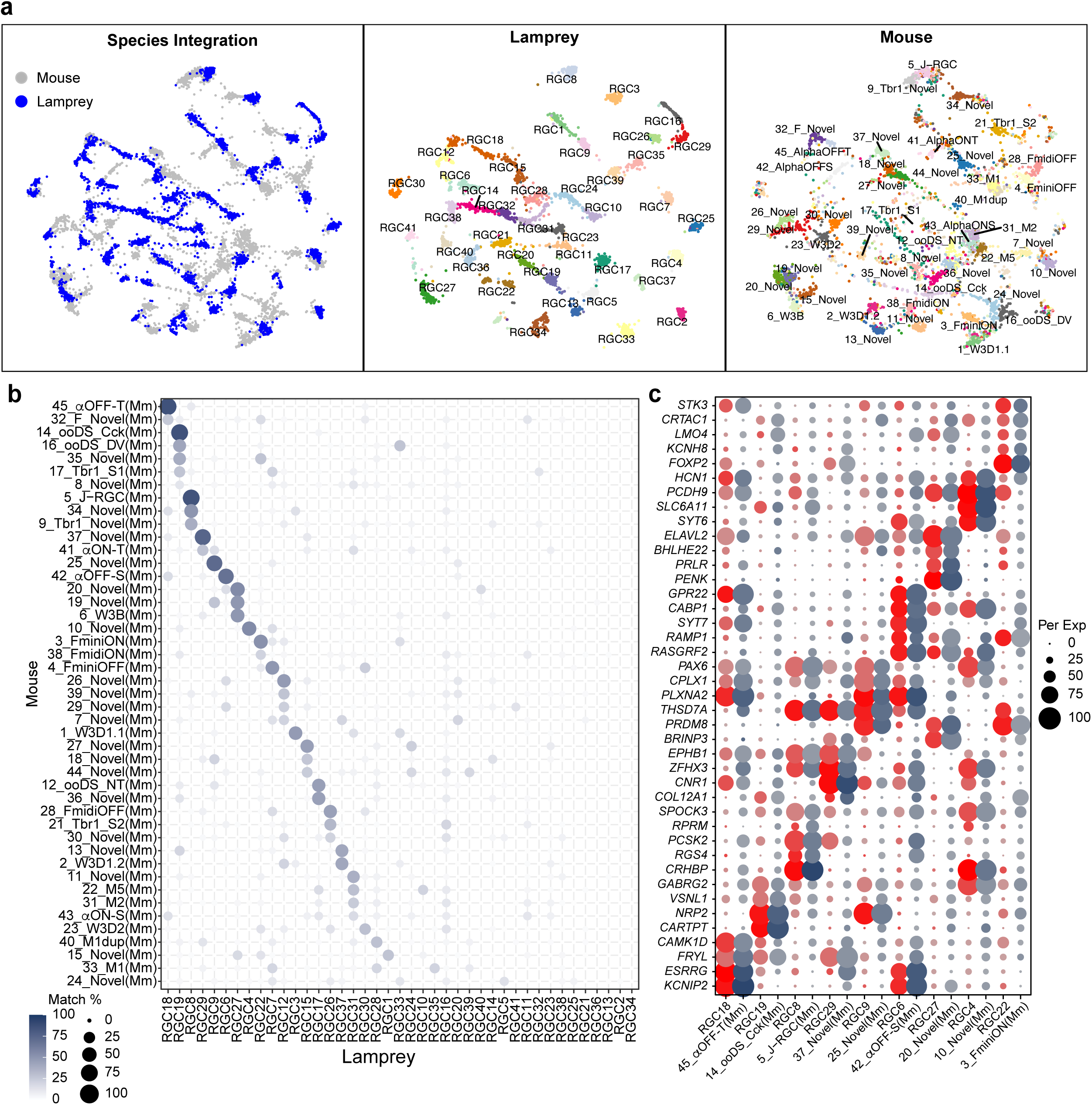
**Comparative analysis between lamprey and mouse RGC types** (a) Integration of lamprey and mouse RGCs visualized with UMAP, with both integrated and species-specific clusters presented in separated UMAP plots. (b) Confusion matrix demonstrating transcriptomic correspondence of RGC types between lamprey and mouse. Lamprey RGCs were used as the training dataset, while mouse RGCs were used as the testing dataset. Mouse RGC types were followed the nomenclature in Ref 44. (c) Dot plot showing expression patterns of conserved markers across corresponding RGC types between lamprey and mouse.

Surprisingly, we didn’t identify a match for ipRGCs between lamprey and chicken or mouse RGCs, suggesting a divergent origin of ipRGCs between jawless and jawed lineages. These results indicate that the level of diversification of RGCs in the lamprey is similar to that in jawed species; however, only a few ganglion-cell types seem to share a common origin with jawed species, suggesting extensive separate evolution in visual processing at the level of the ganglion cell.

### Ancient origin of genetic regulators in differentiating retinal cell classes

The sharing of retinal cell classes across significant phylogenetic distances suggests that gene regulatory programs specifying these classes are ancient. These programs may already have emerged in vertebrate ancestors. To explore these ancient genetic regulators, we applied network-based regulon inference and protein activity analysis to compare the activities of master regulators between the lamprey and a distant mammalian species, the macaque monkey. These master regulators comprise transcriptional factors (TFs) and co-TFs, which drive the expression of cascades of downstream genes to specify the fate of cell classes. Master regulators also include signaling molecules and membrane proteins that influence molecular and physiological features of cell classes.

We used the Algorithm for the Reconstruction of Accurate Cellular Networks implemented with an Adaptive Partitioning strategy (ARACNe-AP) to identify potential master regulators^68^. We then developed the Regulon Structure-based Enrichment Analysis (ROSEA) to infer the protein activities of these master regulators (Fig. 7a) (See Methods)^69^. Through this analytical framework, we identified master regulators in both the lamprey and macaque datasets. Based on the protein activities of these regulators, cells from both lamprey and macaque were grouped into clusters. Each cluster belonged separately to one of the six cell classes (Fig. 7b and 7d). We further identified highly active regulators specific to each cell class (Fig. 7c and 7e). Many of these regulators, such as *ONECUT1*, *VSX2*, and *SOX9*, have been shown to play a crucial role in differentiating retinal cell classes (Fig. 7c and 7e)^46,70–72^. These results demonstrated the precision of protein inferring and clustering and confirmed the specificity of master regulators for individual cell classes in both species.

**Figure 7.**
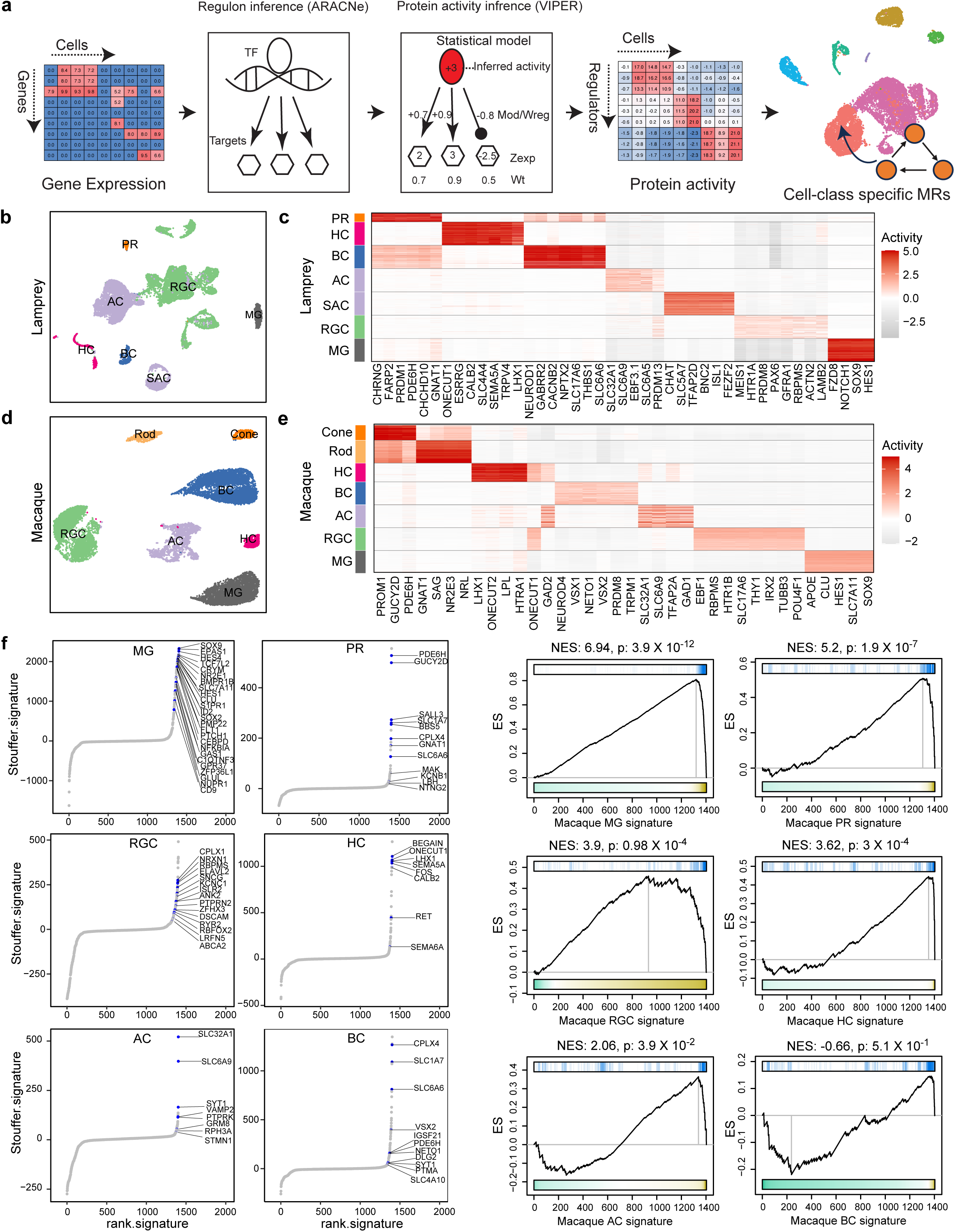
**Identification and comparison of protein activities of class-specific master regulators between lamprey and macaque** (a) Analysis framework for network-based master regulator and protein activity inference. First, the gene regulatory network was reversely engineered from the gene expression profile with ARACNe-AP. ROSEA was then used to infer protein activity from the regulon structure and gene expression status of the targets. Dimension-reduction analysis was conducted, and essential regulators were inferred from protein activity. (b) Lamprey cells were clustered based on protein activities of master regulators and visualized with UMAP. Six cell classes were identified. (c) Heatmap showing the protein activities of class-specific master regulators in lamprey. (d) Macaque cells were clustered based on protein activities of master regulators and visualized with UMAP. Six cell classes were identified. (e) Heatmap showing protein activities of class-specific master regulators in macaque. (f) Waterfall plots depicting inferred top 100 regulators in the lamprey that are also activated in macaque (Z*_stouffer_* > 1.65) as conserved regulators across species. The signature indicates the macaque signature. (g) Comparison of essential regulators for retina cell classes between lamprey and macaque from gene set enrichment analysis. The p values indicate the degree of conservation. NES: normalized enrichment score.

We next compared top-active master regulators present in individual cell classes between lamprey and macaque. First, we identified 1400 regulators shared between the species. Next, we ranked these regulators for each class, considering their integrated protein activities in the macaque dataset, which we referred to as “macaque cell-class signature.” Among top 100 active regulators in each class, we identified multiple common master regulators shared with lamprey cell classes (Fig. 7f). These regulators likely represent some of the most ancient genetic programs, possibly defining original cell classes in our vertebrate ancestors. We then assessed the level of conservation of cell classes between the macaque and lamprey. Our criteria were based on the extent to which a similar set of master regulators is shared between the two species. We employed gene-set enrichment analysis (GSEA) for this purpose and selected the top 50 regulators specific to each lamprey cell class as a gene set of interest. We calculated normalized enrichment scores (NES) and statistical p-values of this gene set referred to the macaque cell-class signature (Fig. 7g) (see Methods). A higher enrichment score indicates a higher level of conservation, signifying that a similar set of master regulators is shared between the two species. Our results showed that, except for BCs, all other cell classes demonstrated significant conservation, as highlighted by their p-values (Fig. 7g). Notably, MG displayed the highest conservation of master regulators among all classes, while BCs had the lowest conservation and are highly different between the two species.

## Discussion

Considerable effort has recently been given to understanding the genetic characterization of different cell types in the nervous system. This study is, to our knowledge, among the first to compare neuronal cell types between lamprey and later emerging vertebrates, groups that diverged in the late Cambrian over 500 million years ago^73^. Our study has shown that certain retinal cell types were clearly established at the time of the separation of cyclostomes from other vertebrate classes, including rod and cone photoreceptors and certain retinal interneurons (such as starburst amacrine cells), which clearly resemble those in later vertebrates. Mechanisms of developmental regulation also appear to have emerged very early in the evolution of the retina.

A previously underestimated aspect of the lamprey retina is its rich variety of cell types, challenging the traditional understanding that early vertebrates possess a simpler nervous system. We identified 74, a number comparable to that in primates^30,74^. The molecular characteristics of these cell types and their parallels to jawed species highlight an impressive correspondence between lamprey and jawed vertebrates. There were, however, also some important differences. Lamprey retina has many fewer amacrine cell types than mammals, and our results suggest that single types of lamprey amacrine cells are related to several types in mammals. Moreover, we found much less correspondence of ganglion cell types between lamprey and mammals than for the other cell classes. Our study has shown that retinal types during evolution were in some cases remarkably stable but in other cases showed considerable divergence, probably reflecting the different behavior and environmental constraints imposed on different species. Our work suggests that evolution proceeded opportunistically, preserving cells and circuits that maintained their usefulness but inventing new mechanisms as these became adaptive.

### The Origin of Duplex Retinal Circuitry

Our work provides new insights into the origin of duplex retinal circuitry^75,76^. A duplex retina denotes a retina comprising rods and cones, which mediate scotopic and photopic vision^32,33^. In the mammalian retina, a designated rod pathway emerged solely for transducing rod signals at absolute threshold^77^. In this pathway, rods primarily contact a single type of rod bipolar cell (RBC), and RBCs don’t synapse directly with RGCs but instead terminate on a unique AC type, the AII AC^78^. Such an adaption is believed to enhance vision in low-light conditions by allowing rod signals to be pooled when few rods absorb photons, and then integrating these signals into cone pathways^79^. The AII ACs then form gap junctions and inhibitory synapses with ON and OFF CBCs, which in turn connect to RGCs^80,81^. This indirect mechanism of rod signaling was once thought to be exclusive to mammals, but recent molecular and connectivity characterizations of the rod pathway in zebrafish have indicated that at least some lower vertebrates may also process rod signals in this fashion^48^.

Our findings suggest that some mechanisms of rod signaling may have originated before the split of jawless vertebrates from other vertebrate ancestors. We have shown that rods are transcriptomically different from cones, and that they use distinct phototransduction gene sets (Fig. 2a, 2b). Our research has also identified RBCs in the lamprey retina (Fig. 3). Lamprey RBCs express the marker genes *PRKCA/PRKACA* and *GRAMD1B* and bear a morphological resemblance to mammalian RBCs (Fig. 3b, 3d). Interestingly, transcriptomic mapping results suggested the presence of two ON-type RBC types in the lamprey, mirroring recent findings of two ON RBC types in zebrafish (Fig. 3f, 3g)^48^. Last, a critical component of the mammalian rod pathway, the AII-AC, is closely aligned to lamprey AC8 (Fig. 5c), though the AC8 cells do not express gap-junction genes. It is therefore possible that a rod pathway like that in mammals exists in lamprey and may have originated in common vertebrate ancestors of jawed and jawless vertebrates.

It is however also possible that lamprey have a more primitive pathway for rod signaling, instead of or in addition to the AII-amacrine pathway now utilized by mammals. Rods evolved from cones and may have initially utilized bipolar cells of both ON and OFF types similar to ON and OFF cone bipolar cells, directly contacting ganglion cells. It may be significant in this regard that lamprey have been shown to have a purely rod bipolar cell which is OFF or hyperpolarizing^49^. Our observations also indicate that there is an OFF-BC type, BC7, which expresses the RBC marker *GRAMD1B* and may be the OFF RBC previously identified^49^. Moreover, OFF rod bipolar cells have been detected in the retina of amphibians^82–84^. These findings suggest that ON and OFF rod bipolar cells were present in lamprey progenitors before the split of cyclostomes from other vertebrate lineages. OFF rod bipolar cells may then have slowly disappeared in some lineages during evolution, as this more primitive direct bipolar-to-ganglion-cell pathway was replaced by the AII pathway now found in mammals and some other vertebrate species^76^. Future investigation into the morphology, physiology, and connectivity of lamprey bipolar and amacrine cells will be crucial for exploring these hypotheses and determining how rod pathways in lamprey resemble or differ from those in other vertebrate species.

### ON and OFF Pathways

Another notable feature of the lamprey retina is the presence of ON and OFF pathways, which discern the increment and decrement of light. The distinction between ON and OFF begins at synaptic connections between photoreceptors and bipolar cells in the outer plexiform layer (Fig. 1b), where BCs can be categorized into ON and OFF subclasses^85^. This functional differentiation is due to opposing responses to glutamate. In the dark, glutamate released by PRs depolarizes OFF BCs by activating their ionotropic glutamate receptors^43^. In contrast, photoreceptors hyperpolarize ON BCs via metabotropic glutamate receptors, which subsequently close constitutively active non-selective cation channels now known to be transient-receptor-potential melastatin (TRPM) channels^86^. In mouse retina, OFF BCs can express the kainate receptor GRIK1, while ON BCs typically express GRM6 and TRPM1^87^. A previous study using functional recording demonstrated the presence of ON and OFF BC types in the lamprey^88^. This study showed that the light responses of ON BC are sensitive to AP4, suggesting lamprey ON BCs express group III metabotropic glutamate receptors. Our results align with this functional result and have further revealed that lamprey ON BCs express GRM8, a different group III metabotropic glutamate receptor, rather than GRM6 (Fig. 3b, Supplementary 4a). Furthermore, it seems that GRM8 associates with TRPM3 instead of TRPM1 in lamprey ON BCs (Fig. 3b, Supplementary Fig. 4a). Additionally, lamprey OFF BCs express GRIK2 instead of GRIK1. Thus, our results confirm that BCs in the lamprey retina differentiate into ON and OFF subclasses, and their distinct glutamate receptor usage suggests that some aspects of transmission between photoreceptors and bipolar cells were more variable in primordial vertebrates than in present-day jawed species.

An ON and OFF distinction extends into the inner plexiform layer (IPL) via specific synaptic interactions between presynaptic ON/OFF BCs and postsynaptic ON/OFF types of amacrine cells or retinal ganglion cells. Notably, starburst amacrine cells (SACs) are among the earliest cell types to stratify their dendrites in the IPL and are classified into ON and OFF types^89^. ON and OFF SACs have been shown to offer scaffoldings for the innervation of respective BC types and play a pivotal role in organizing the ON and OFF circuitry in the mouse retina^89,90^. Our study not only detected ON and OFF SACs in the lamprey retina but also observed similar gene expression patterns to those in mice (Fig. 4). Specifically, *FEZF2* is expressed by ON SACs, echoing the expression pattern of its mouse paralog, *Fezf1*, which determines the fate choice of ON versus OFF SACs^57^. Intriguingly, transmembrane proteins, such as TENM3, are also expressed differentially between lamprey ON and OFF SACs (Fig. 4l). These results suggested that a potentially analogous molecular mechanism governs ON and OFF laminations in the lamprey IPL. Additionally, *MEGF10*, which regulates the spatial arrangement of SACs in the mouse retina, is also present in lamprey SACs^55^. Altogether, the existence and division of SACs into ON and OFF subgroups seems to be a primitive and conserved trait across all vertebrates.

### Unique Features of the Lamprey Retina and Evolutionary Modifications

The lamprey retina possesses several unique features. First, there are four types of horizontal cells (HCs) identified by our study in the sea lamprey retina, even though there is only a single spectral class of cone in this species^91,92^. Typically, the presence of multiple types of HCs correlates with diverse types of cones, as observed in the retinas of zebrafish, chickens, and macaques^93–95^. Similarly, the mouse retina has only one cone type, and it has one HC type^96,97^. There are however other lamprey species with several chromatic classes of cones, which may require a larger diversity of horizontal cell types^98^. The circuit connectivity and physiological function of these diverse HC types in lamprey merits future exploration.

Second, all lamprey ACs express both GABAergic and glycinergic transporters. Furthermore, each lamprey AC type corresponds to multiple types of mouse or chicken ACs. These results may be explained by the theory of “apomere” evolution of new cell types via module divergence. In this theory, ancestral AC types could be multifunctional with multiple modules, and the diversification of AC types in jawed species may have occurred through the segregation of functions or modules into distinct sister AC varieties^26,99^.

Lastly, RGCs in the lamprey retina feature the greatest diversity among all the cell classes, with 41 distinct types. This count is comparable to the number of RGC types in mouse or chicken retinas. However, there is a stark contrast in terms of conserved RGC types between lamprey and other species. These results reflect a major difference in the localization of the somatic and axonal layer of lamprey RGCs compared to jawed species (Fig. 1b). Jones et al.^60^ and Fletcher et al.^61^ identified six morphological RGC groups in lampreys: two inner ganglion cell (IGC) groups, IGCa and IGCb; three outer ganglion cell (OGC) groups, OGCa, OGCb, and OGCc; one bipolar ganglion cell group, BPGC. Of these, four (IGCa, IGCb, OGCa, OGCb) are homologous to RGCs in other vertebrates. These cells stratify their dendrites in the IPL and project to the tectum, directing eye, head, and body movements. Intriguingly, a couple of RGC types in the lamprey align with several direction-selective ganglion cells (DSGCs) including ON-OFF DGSCs and J-RGCs, and also local motion detector W3 cells in the mouse retina^62,64,67^. These results support the role of lamprey RGCs in mediating phototaxis ^100^. Moreover, since DSGCs connect with SACs, these correspondences may reflect conserved synaptic partnership with SACs, and direction-selective circuits for motion detection may have been one of the earliest design features of the vertebrate camera eye, enabling gaze stabilization and object tracking^53^. The remaining two subtypes (BPGC and OGCc) extend their dendrites into the OPL, directly contacting PRs and projecting to the pretectum, potentially regulating dorsal light and visual-escape responses. Thus, we might infer that some RGC types were peculiar to jawless species and could have been lost in the evolutionary transition to jawed species. Additionally, the lack of correspondence of ipRGCs between lamprey and macaque suggests a distinct function of ipRGCs in the two species^101^.

In this study, we also emphasize a method of inferring regulatory protein activity from scRNA-seq datasets. While scRNA-seq provides a genome-wide profiling of gene expression in individual cells, the expression level of a gene may not always correspond to its protein activity due to post-translational modification^102^. Moreover, transcription factors involved in cell-type specification during development may have reduced expression in adult cells. Given the pleiotropic nature of transcriptional regulation and the evolutionary changes in the co-regulatory complex of transcription factors^26^, we used network-based methods. These methods consider the regulatory structure–the relationship between gene expression of regulators and their targets, as well as the expression status of target genes to infer the activity of master regulators. With this approach, our method identified ancient regulators that might have originated in common vertebrate ancestors. Our study also suggests that the BC class specification may be the least conserved among all retinal classes between lamprey and macaque. This finding could highlight a distinction in the generation of BCs between jawless and jawed species. In jawed species, BCs are the last cell class to be generated and to mature. In contrast, BCs are evident in larval lamprey prior to the generation of HCs and ACs^12^.

### Limitations of our study

Despite the great diversity of cell types identified in this study, additional cell types could likely be revealed with the sequencing of more cells, including additional types among bipolar cells. Our characterization of cell types in the lamprey is primarily based on transcriptomic distinctions. Many of the cell types identified in this manner warrant future histological and function validation. In particular, the identification of a wide range of RGC types, along with future clarification of their localization, dendritic projections, and brain innervation, will provide important insight into understanding the diversification of RGC types in vertebrates.

From the improved gene annotation of the lamprey genome, we were able to project the best corresponding cell types between lamprey and jawed species using integration and transcriptomic mapping methods. The purpose of this survey is to provide a preliminary examination of ancestral cell types that might have been emerged before the divergence of the jawless and jawed vertebrate lineages.

## Acknowledgements

This work was supported by a career development award from Research to Prevent Blindness, a career starter award from Knights Templar Foundation, a Klingenstein-Simons Neuroscience Fellowship (Y.R.P.), NEI grants R01 EY001844 (G.L.F.) and R01 EY029817 (A.P.S.), a grant from the Great Lakes Fishery Commission (G.L.F.), and an unrestricted grant to the Department of Ophthalmology from Research to Prevent Blindness, Inc., and Core Grant P30 EY00331 to the Jules Stein Eye Institute.

## Author Contributions

Y.R.P. conceived and supervised the study. J.W., and L.Z. performed the computational analysis with the assistant from J.S.; Y.R.P. performed TruSeq and scRNA-seq experiments. A.P conducted the histology and in situ experiments with the assistant from M.C.; A.M. collected and dissected tissues under the supervision of G.L.F. and A.P.S.; Y.R.P. and G.L.F. wrote the paper with input from other authors.

## Methods

### Tissue procurement

Lamprey tissue collection was carried out in accordance with the recommendations of the Guide for the Care and Use of Laboratory Animals of the National Institutes of Health, USA, and was approved by the University of California, Los Angeles, Animal Research Committee. Sea lamprey, *Petromyzon marinus* Linnaeus 1758, were provided by the Hammond Bay Biological Station of the United States Geological Survey (USGS), Millersburg, MI, USA. They were kept in a large fresh-water aquarium at 4°C on a 12 h:12 h light:dark cycle. Lampreys were deeply anesthetized with 400 mg/L tricaine methanesulfonate (MS-222; E10521, Sigma–Aldrich, St. Louis, MO) and decapitated. After dissecting out eyes, the anterior chamber and the vitreous were removed by rapid hemisection. The posterior eyecup was immersed in room-temperature Ames’ medium (Sigma), equilibrated with 95% O2/5% CO2 for at least 20 mins) for cell dissociation, or immediately fixed with 4% PFA for immunohistochemical experiments or fluorescence in situ hybridization.

### Fluorescence In Situ Hybridization (FISH) validations

After isolation of the tissue, the posterior poles containing the retina were fixed with 4% PFA for 1hr at 4°C, rinsed with PBS and immersed in 30% sucrose at 4°C overnight, and then embedded with Tissue Freezing Medium. The tissue was sectioned at 20-μm thickness and stored at -80°C for long-term storage. Fluorescence in situ probes against specific lamprey genes were generated with previously described methods (Peng 2019). Briefly, total RNA was extracted from lamprey retinas and converted to cDNA libraries through reverse transcription with the AzuraQuant cDNA synthesis kit. Antisense probe templates for individual target genes were PCR-amplified from the cDNA libraries with a reverse primer having a T7 sequence adaptor to permit in vitro transcription. DIG rUTP (Roche) and Fluorescein rUTP (Roche) were used to synthesize probes for single or double FISH experiments. Retinal sections were thawed, treated with 1.5 mg/mL of proteinase K (NEB), post-fixed with PFA, and deacetylated with acetic anhydride. After blocking, the retinal sections were incubated with probes overnight. Probe detection was performed with anti-DIG HRP (1:1000) and anti-Fluorescein POD (1:1000) followed by tyramide amplification^103^.

### Immunohistochemistry

Tissue was fixed and prepared as described above. Antibodies used were as follows: anti-Calbindin (1:2000, Swant); mouse and rabbit anti-PKCa (1:2000, Abcam; 1:2000, sigma). Nuclei were labeled with DAPI (1:1000, Invitrogen). Secondary antibodies were conjugated to Alexa Fluor 488 (Invitrogen) and used at 1:1000. Sections were mounted in ProLong Gold Antifade (Invitrogen).

### Image Acquisition, Processing, and Analysis

Images were acquired on an Olympus FluoViewTM FV1000 confocal microscope with 405, 488, and 599 lasers and scanned with a 40X or 60X oil objective at a resolution of 1024×1024 pixels, a step size of 1 μm, and an 80 mm pinhole size. Maximum intensity projections were generated with ImageJ^104^ (NIH) software, and brightness and contrast adjustment were made with Adobe Photoshop.

### Construction of a retina-specific transcriptome for *Petromyzon marinus*

We obtained high quality total RNA from lamprey retinal tissue [RNA Integrity Number (RIN) score: 9.8], and prepared strand-specific libraries with the TruSeq strand-specific Total RNA kit (Illumina Inc.), which was sequenced on the NextSeq 500 system to obtain 50 million 100bp paired end reads. We merged TruSeq bam files of samples S1 and S2 by “samtools merge”^105^ and generated a merged fastq file with “samtools bam2fq”. We updated the gtf file of NCBI genome assembly kPetMar1.pri (https://www.ncbi.nlm.nih.gov/datasets/genome/GCF_010993605.1/) of sea lamprey (*Petromyzon marinus*) according to the following steps: (1) We used HISAT2^106^ for reads alignment. We used the hisat2_extract_splice_sites.py and hisat2_extract_exons.py scripts to extract spicing sites and exons. We then used “hisat2-build” to create hisat2 index. The final bam file was then generated by hisat2. (2) We used StringTie^107^ for transcript assembly and quantification. The gtf file was updated by stringtie by using the HISAT2 bam file. The stringtie gtf and the reference gtf were compared with gffcompare^108^ and merged with stringtie–merge. Transcript abundances were estimated with stringtie (with arguments -e -B). After the update, the NCBI+TruSeq reference transcriptome had 12,061 genes updated by stringtie with MSTRG Number. We used the updated transcriptome for aligning scRNA-seq reads (see below).

The NCBI reference genome of the sea lamprey (kPetMar1.pri) has over 60% of its genes assigned with a LOC number, which means that these genes had uncertain functions. To facilitate cell type classification with canonical retina markers, we further annotated these genes based on the identification of their protein products. We labelled genes sharing the same protein by adding a suffix to the gene name (Supplementary Table 1). We used upper case for all the lamprey gene symbols. With this method, we improved the proportion of annotated genes in the reference transcriptome from 36.22% to 79.44% (Supplementary Fig. 1c).

### Single cell RNA-sequencing Library preparation

The lamprey retinas were dissected out and dissociated into single-cell suspension for single cell RNA sequencing (scRNA-seq, 10x Genomics). Libraries were sequenced on the Illumina NovaSeq S4 platform.

## Single-cell RNA-seq Data Analysis

### Read alignment and generation of count matrices

We used the updated “NCBI+TruSeq” transcriptome to align scRNA-seq reads with Cellranger (10x Genomics, version 7.0.1)^109^. We first constructed a reference transcriptome with “cellranger mkref”. We then mapped scRNA-seq reads with “cellranger count”. The resulting count matrices have gene IDs from the TrueSeq update with MSTRG number or LOC number. To facilitate downstream analysis, we updated count matrices by replacing gene IDs, which are either MSTRG or LOC number, with annotated gene symbols following Supplementary Table 1.

### Data pre-processing, normalization, dimension reduction, and clustering analysis

The raw filtered count matrices were used for further analysis with the Seurat R package (version 4.3.0)^110^. We first removed low-quality cells and putative doublets by examining the distributions of numbers of expressed genes, RNA counts, and percentages of mitochondrial genes detected in individual cells. Data normalization and identification of highly variable genes (HVGs) were performed with the “SCTransform” function. A Gamma-Poisson generalized linear regression (glmGamPoi) model was used for the normalization^111^. We then performed principal component analysis (PCA) with HVGs and further eliminated batch effects with Harmony^112^. After Harmony correction of the top 50 PCs, a K-nearest neighbor (KNN) graph was constructed with Harmony components, and cells were clustered with the Louvain algorithm. The uniform manifold approximation and projection (UMAP) dimension reduction were used for visualization.

### Lamprey retinal cell class annotation

We annotated lamprey retinal cell clusters into distinct classes with the reference-based method SingleR^113^. First, we generated the reference dataset by selecting a small subset of clusters, each of which showed strong specific expression of canonical marker genes for certain cell classes. The remaining clusters were designated as a query dataset. Then we used the log2(CPM+1) transformed data as input for SingleR to make prediction. We assessed the annotation by conducting a detailed examination of class-specific markers for each cluster. We also evaluated the clustering patterns of these classes in the UMAP space generated by changing the number of Harmony components used. We observed that, when using the top 7 Harmony components, cells from the same class predominantly clustered together in the UMAP, indicating the accuracy of the final annotation.

### Cell type classification within each class

To resolve cellular heterogeneity at a higher resolution, we divided cells into individual classes and further clustered cell types within each class. We performed data normalization, PCA analysis, batch correlation with Harmony, and clustering. After clustering, we examined the quality of clusters by evaluating their numbers of expressed genes, RNA counts, percentages of mitochondrial genes, and the expression level of cell-class marker genes. Low-quality cells and clusters with contaminated cells were removed, and a repeat round of clustering analysis was performed until every cluster containing high-quality cells.

### Cross-species integration of the retinal cell classes

The gene names of mice and chicken were converted into human symbols with orthologous relationships given in Ensembl (http://www.ensembl.org/). The genome versions mouse GRCm39 and chicken bGalGal1.mat.broiler.GRCg7b, and human GRCh38.p14 were used.

We used canonical correlation integration (CCA) in Seurat (v4.3.0)^110,114^ for cross-species integration. First, we normalized samples from each species by fitting the Gamma-Poisson generalized linear regression model implemented in the “SCTransform” function. We then selected conserved HVGs with the “SelectIntegrationFeature” function. After finding anchors with the “FindIntegrationAnchors” function with parameter dims=1:50, the datasets were integrated by using “IntegrateData” with parameter dims=1:50. After integrating data, we performed dimension reduction and clustering analysis with the top 50 PCs. To ensure a balanced representation across the diverse cell types within both AC and RGC classes, we downsampled these groups. For lamprey and mouse RGC integration, a slight modification of the selecting integration feature step was applied. First, we identified mouse and lamprey HVGs by running the SelectIntegrationFeature” function for each species. Then, we identified the shared conserved HVGs for these two species by intersecting the mouse HVGs and lamprey HVGs. After that, the datasets from the two species were integrated with “IntegratedData,” followed by dimension reduction and clustering.

### XGBoost mapping

To identify the correspondence of cell types across species, we performed a supervised multi-class classification with XGBoost^115^ (R package version), a scalable machine learning algorithm that uses the gradient tree boosting technique. Implementation comprised two steps: First, a predictive model was built from a training dataset from one species with labeled cell types; Second, the model was used for testing the dataset from another species to predict the corresponding cell-type label. The details of analysis are outlined as follows. We first identified the common HVGs between species in the integrated assay with the “SelectIntegrationFeatures” function in Seurat, and we used these genes as features in XGBoost. We constructed a predictive model from 67% of the cells from each cluster in the training dataset (with a maximum of 300 cells per cluster if the cluster contained more cells), and we used the remaining cells to evaluate the model’s prediction performance. After the model achieved high performance, we applied the testing dataset to it. Finally, we used a confusion matrix to demonstrate the correspondence of cell types between species. A higher matched percentage indicates a higher similarity between cell types.

### Detection of conserved marker across species and across samples

First, the differentially expressed genes (DEGs) within individual species or samples were identified with the Wilcoxon test, as implemented by the “FindAllMarkers” function in Seurat. To eliminate batch effects from different species or samples, we selected the conserved makers with the Stouffer’s integration method^116^. The individual p-values from the DEG tests were transformed into Z scores. Subsequently, these Z scores were combined by using the formula below:

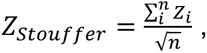

where *i* is the species/sample index.

Furthermore, the Stouffer-integrated Z scores were converted into p-values under the assumption of a standard normal distribution, and they were then adjusted with the False Discovery Rate (FDR) method. To avoid infinity of for Z scores, p values equal to 0 were replaced by the smallest positive double-precision number in R during conversion.

### The construction of phylogenetic tree (dendrogram) among clusters

The correlation distance is defined as

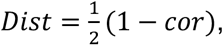

where *Dist* refers to distance and *cor* refers to the Pearson correlation. To generate the dendrogram, the hclust function in the stats R package with centroid agglomeration was used. Bootstrap was performed by using ape^117^ with 100 replicates generated. The CompexHeatmap^118^ R package was used for the heatmap visualization of the correlation coefficients.

### TF, coTF and surface/surface signaling gene assignment in macaque

We used the Gene Ontology (GO) annotations for human transcription factor^119^, transcription cofactor (coTF), and surface/surface signaling genes to assign functions to macaque genes. The list of potential regulators comprises 1,430 transcription factors (TFs), 2744 coTFs, and 2717 surface and surface signaling genes. The GO terms associated with these genes are listed in the following. TFs: “DNA-binding transcription factor activity” (GO:0003700)^120^. coTF (co-Transcription Factors and chromatin remodeling enzymes): “transcription coregulator activity” (GO:0003712); “DNA binding” (GO:0003677); “transcription factor binding” (GO:0008134); “regulation of transcription; DNA-templated” (GO:0006355); “histone binding” (GO:0042393), and “chromatin organization” (GO:0000790) [10]. Surface proteins and surface signaling proteins: “surface proteins” (GO:0009986); cell-cell signaling “GO:0007267” and cell surface receptor signaling pathway “GO:000716”.

### TF, coTF and surface/surface signaling gene assignment in lamprey

We determined orthologous relationships between lamprey and human genes from two different sources. We identified one-to-one (1-1) mapping of orthologous genes from the NCBI annotation, wherein the corresponding orthologous genes share the same gene name. We then determined orthologous genes with OrthoFinder^121^, a tool that uses a tree-based phylogenetic approach. For this procedure, peptide sequences were retrieved from Ensembl, and a customized R script was used to process redundant peptides to ensure that only the longest peptide for each gene was selected for further analysis. From the OrthoFinder results, we identified orthologous genes, including one-to-many, many-to-one, and many-to-many matched types within the assigned OrthoGroups. Finally, the TFs, coTFs, and surface/surface signaling genes in lamprey were annotated based on the GO terms for human ortholog genes.

### Reverse engineering the gene regulatory networks

Regulatory networks were reverse-engineered with ARACNe-AP^68^. ARACNe-AP was run with default settings: bootstrap n=200, mutual information threshold “P<10^-8^” and DPI (data processing inequality; enable=yes). We ran ARACNe-AP on a subset of the single-cell data. To handle the imbalance of the dataset, we used the downsampling technique to ensure that a substantial number of cells were included from classes with a small cell population.

We generated the network for each sample to avoid a batch effect. The TF, coTF and surface/signaling proteins were used independently to generate their networks. These tree networks were merged to form a sample-specific network. Further, we used VIPER^69^ R package to generate regulons from the networks. We pruned the regulons up to size 50 (up to 50 targets for each candidate regulator) and removed the regulon whose size was smaller than 20 for downstream protein activity analysis.

### Protein activity inference and master regulator inference

Protein activities were inferred with Regulon Structure-based Enrichment Analysis (ROSEA, https://github.com/JunqiangWang/Rosea). ROSEA is an algorithm to perform regulon structure-based enrichment analysis according to a probabilistic model. We inferred protein activities for each sample from pruned sample-specific networks as input. An average method was applied to integrate activity scores inferred from different sample-specific networks.

We performed dimension reduction and clustering analysis using the protein activities. We calculated standard deviations of protein activities and selected features by choosing the top 700 proteins with high standard deviations for PCA analysis. Then we performed PCA, batch correction with Harmony, and UMAP dimension reduction based on the top 50 PCs. Further, we performed differential protein activity analysis with the Student’s T test to infer the class-specific regulators. Customized R scripts were used to perform the analysis by implementing functions in Seurat.

### GSEA analysis

Gene-set enrichment analysis was performed for estimating the conservation of essential regulators across species. Class-specific signatures were generated by using Stouffer’s method to integrate the protein’s activity score (Z score) of individual cells within each class. The top 50 highly activated proteins in the signature for each lamprey retina class were used to query whether these proteins are overrepresented in macaque class-specific ranked signatures.

### Data Availability

The retinal lamprey scRNA-Seq data has been deposited to the Gene Expression Omnibus.

### Code Availability

Codes used in this paper are available at https://github.com/PengYRLab/LampreyRetinalCellAtlas.

