## Supplementary Figures for "Molecular Characterization of the Sea Lamprey Retina Illuminates the Evolutionary Origin of Retinal Cell Types"

**Supplementary Information**

**Supplementary Figure Legends**

**Supplementary Figure 1. Assembly and annotation of retina-specific transcriptome.**
(a) Bar plots showing mapping percentages of scRNA-seq reads to "Transcriptome," "Exon," and "Genome" from two biological replicates (S1 and S2) with Ensembl (Pmarinus_7.0), NCBI (kPetMar1.pri), or the updated NCBI+TruSeq transcriptome reference.
(b) Improved gene-body definition for the *red-opsin* gene in the NCBI+TruSeq transcriptome. The alignment of scRNA-seq (top panel) and TruSeq (bottom panel) reads to the "*red-opsin"* locus was visualized with the Integrated Genomics Viewer (IGV). A red arrowhead indicates a newly identified exon region of the *red-opsin* gene
(c) Pie charts showing the proportions of genes annotated with LOC numbers, MSTRG numbers, or gene symbols in the raw count matrices. Three different gtf files are compared: the NCBI gtf, the NCBI+TruSeq gtf, and the NCBI+TruSeq gtf with updated gene annotation.

**Supplementary Figure 2. Quality metrics of lamprey scRNA-seq data and cell type hierarchical relationships.**
(a) UMAP visualization of all lamprey cells, as depicted in Figure 1C, but here colored by distinct replicates to demonstrate lack of bias associated with different replicates.
(b) Violin plots showing distributions of the number of expressed genes (nFeature_RNA), RNA counts, and percentages of mitochondrial genes (percent.mt) detected in each replicate.
(c) Bar plot displaying fractions of cells from each replicate across individual cell types.
(d) Hierarchical clustering and heatmap showing the Pearson correlation coefficients calculated for each pair of cell types, using the top 3,000 highly variable genes.

**Supplementary Figure 3. Fluorescence in situ validation of marker genes for lamprey PR types**
(a) Fluorescence in situ hybridization (FISH) validations confirming exclusive expression patterns of *red-opsin* and *GNAT1* in PR2 and PR1. *Red-opsin* (green) is expressed by PRs with long outer segments (LOS, indicated by an arrow); while *GNAT1* (magenta) is expressed by PRs with short outer segments (SOS, indicated by an arrowhead). The LOS and SOS structures are more clearly visualized in the differential interference contrast (DIC) image.
(b) FISH validation showing the exclusive expression of *Rhodopsin* (*RHO*) and *GNAT2* in PR1 and PR2. *RHO* (green) is expressed by PRs with short outer segments (SOS, arrowhead); while *GNAT2* (magenta) is expressed by PRs with long outer segments (LOS, arrow).
Nuclei are stained with DAPI. Scale bar, 20 $\mu$m.

**Supplementary Figure 4. Expression of ion channels and the hierarchical relationship among BC types.**
(a) Dot plot showing the expression patterns of genes from the TRPM, GRM, and GRIK families across BC types.
(b) Hierarchical clustering and heatmap displaying Pearson correlation coefficients calculated between each pair of BC types using the top 3000 highly variable genes. The dendrogram on the left shows their hierarchical relationships, constructed from agglomerative hierarchical clustering based on correlation distance.

**Supplementary Figure 5. The comparison of the AC types between lamprey and chicken.**
(a) Integration of lamprey and mouse RGCs visualized with UMAP, with both integrated and species-specific clusters presented in separate UMAP plots.

(b) Confusion matrix demonstrating transcriptomic correspondence of AC types between lamprey and chicken (Gg). Lamprey ACs were used as the training dataset, while chicken ACs were used as the testing dataset. Dot size shows the percentage of ACs in each chicken AC type matched to each lamprey AC type.
(c) Dot plot showing the expression patterns of conserved markers across corresponding AC types in lamprey and chicken (Gg).

**Supplementary Figure 6. Expression patterns of marker genes for RGC subgroups and types.**
Dendrogram at the top gives the hierarchical relationships among RGC types, constructed from agglomerative hierarchical clustering based on correlation distance. RGC types are grouped into seven subgroups based on Person correlation coefficients calculated between each pair of RGC types from the top 3,000 highly variable genes. These seven RGC subgroups are visually distinguished by different colors and enclosed within shadow boxes. Subgroup-specific markers are shown in the middle dot plot, while the expression patterns of markers for individual RGC types are presented in the bottom dot plot.

### Supplementary Figure 1

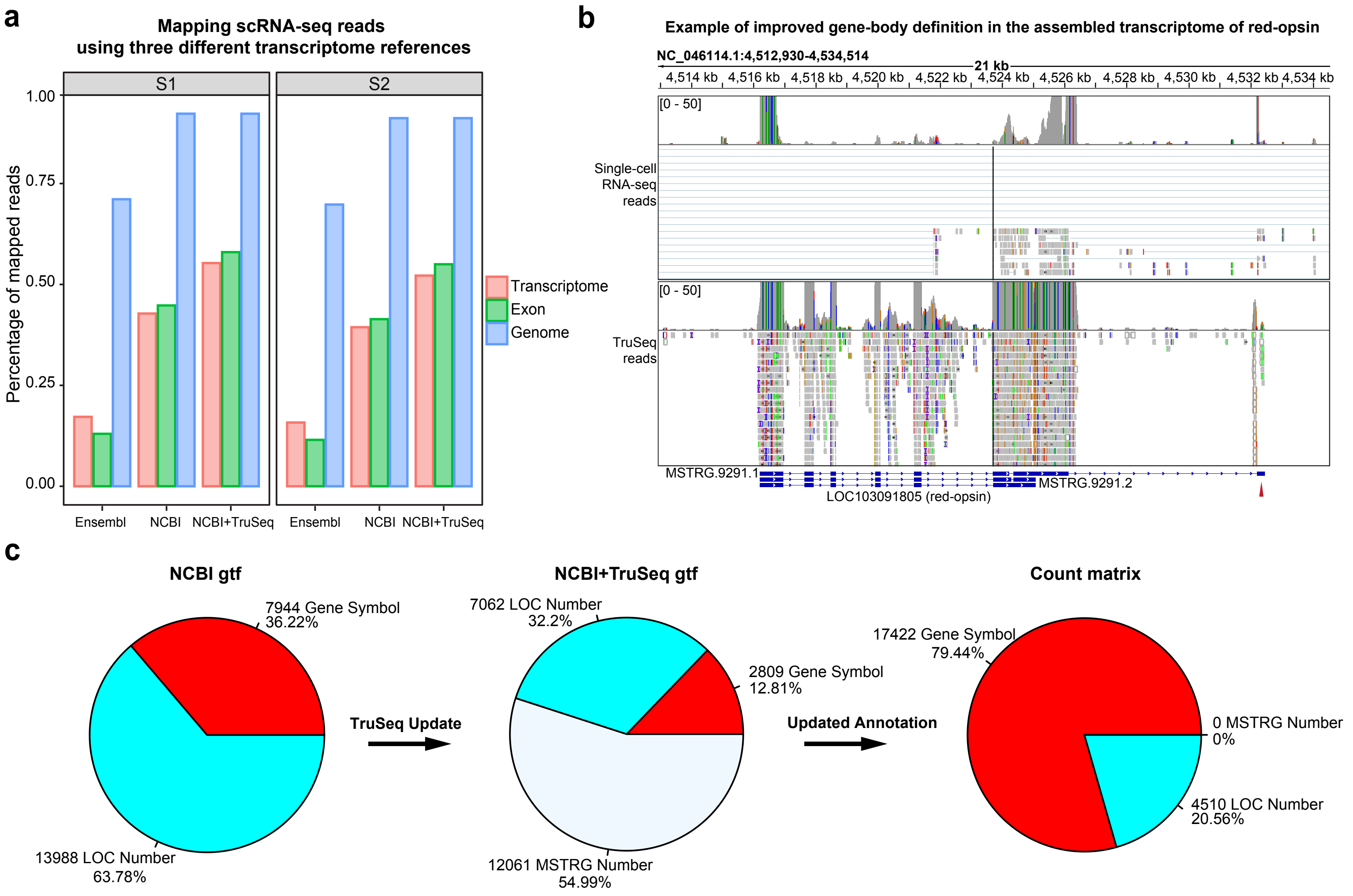

### Supplementary Figure 2

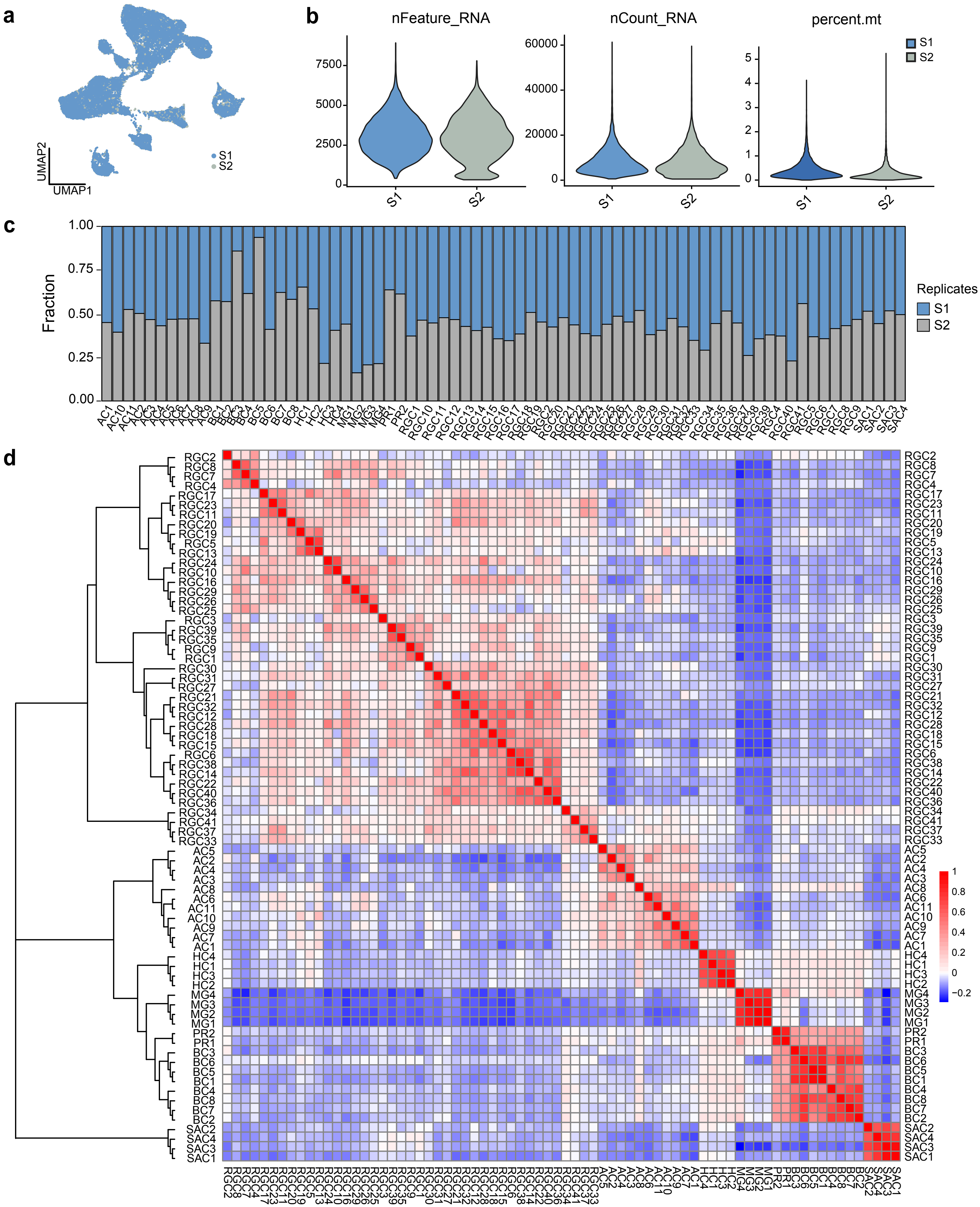

**Supplementary Figure 3**

**a**

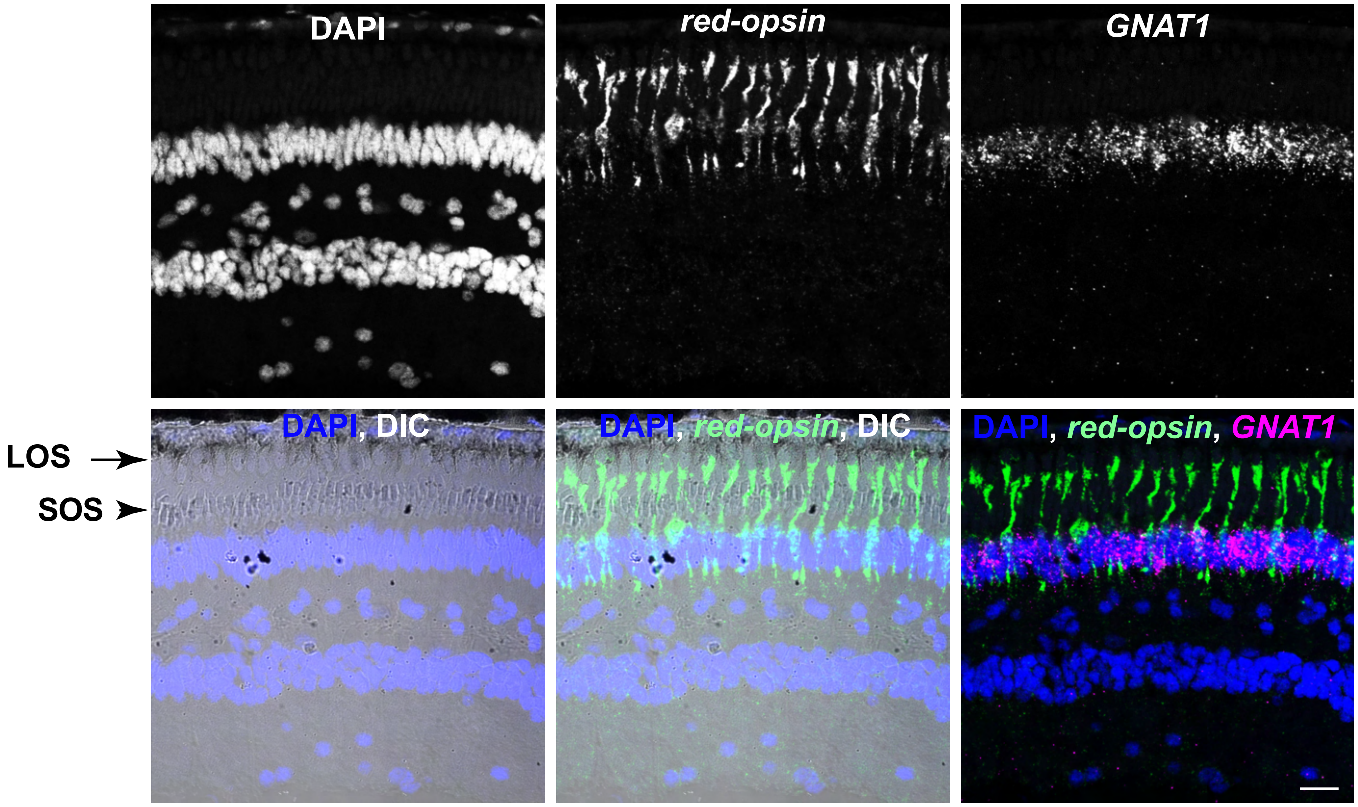


**b**

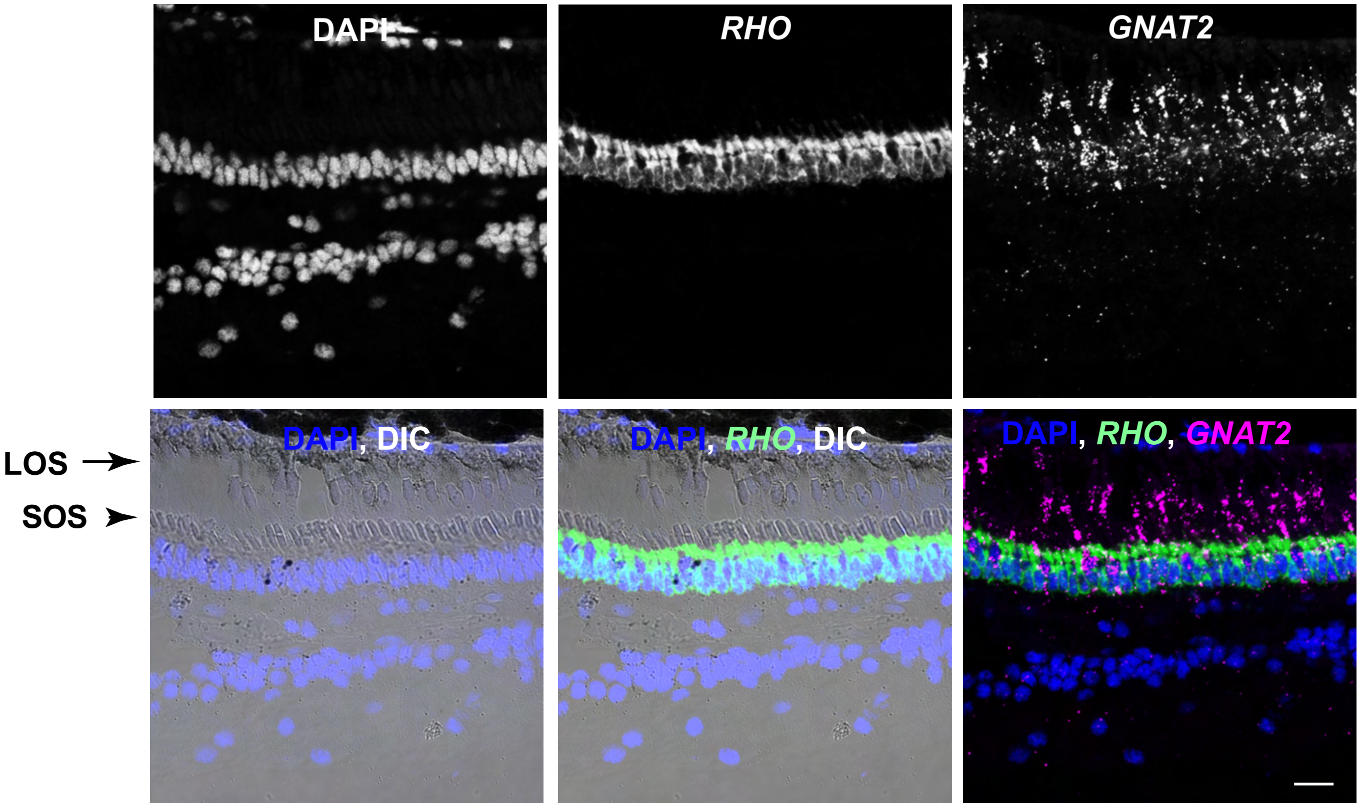

### Supplementary Figure 4

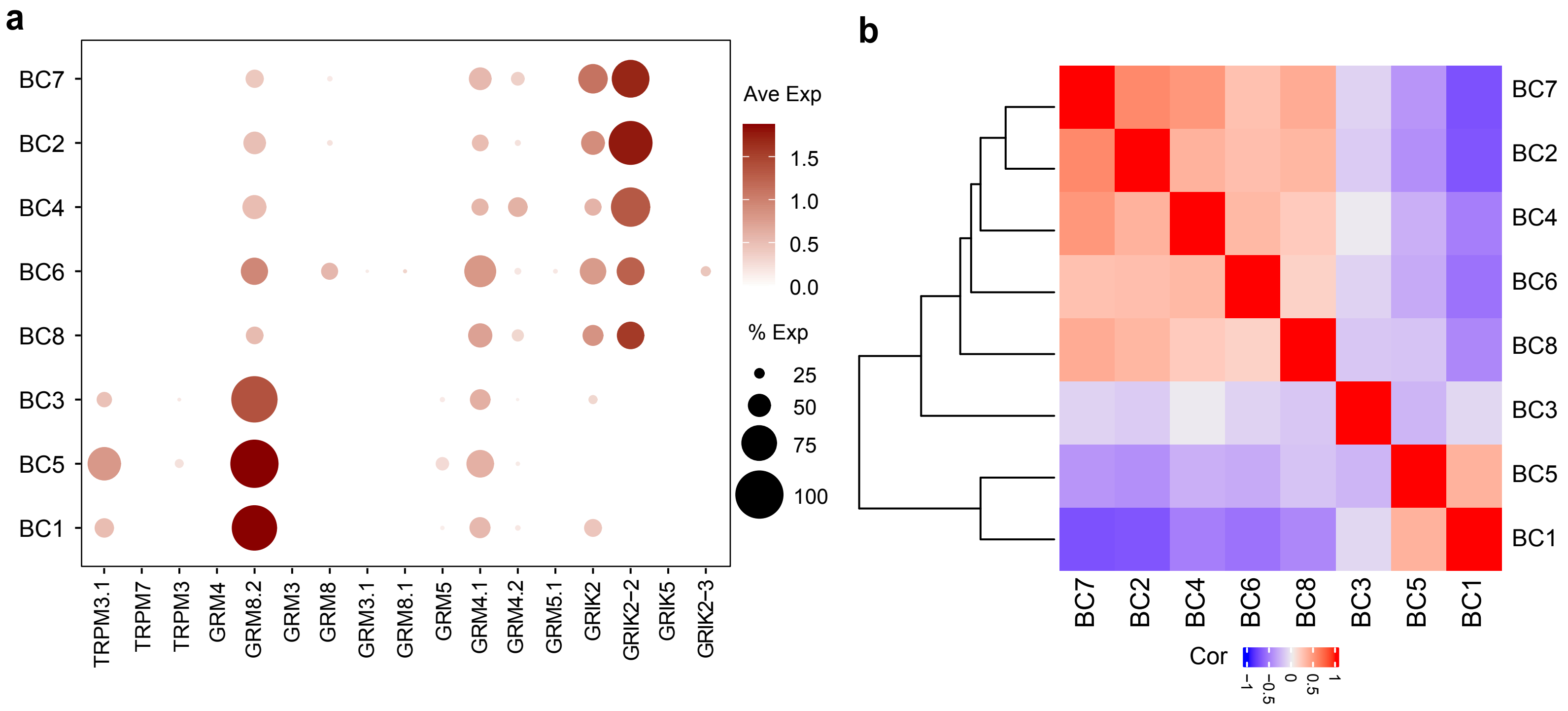

### Supplementary Figure 5

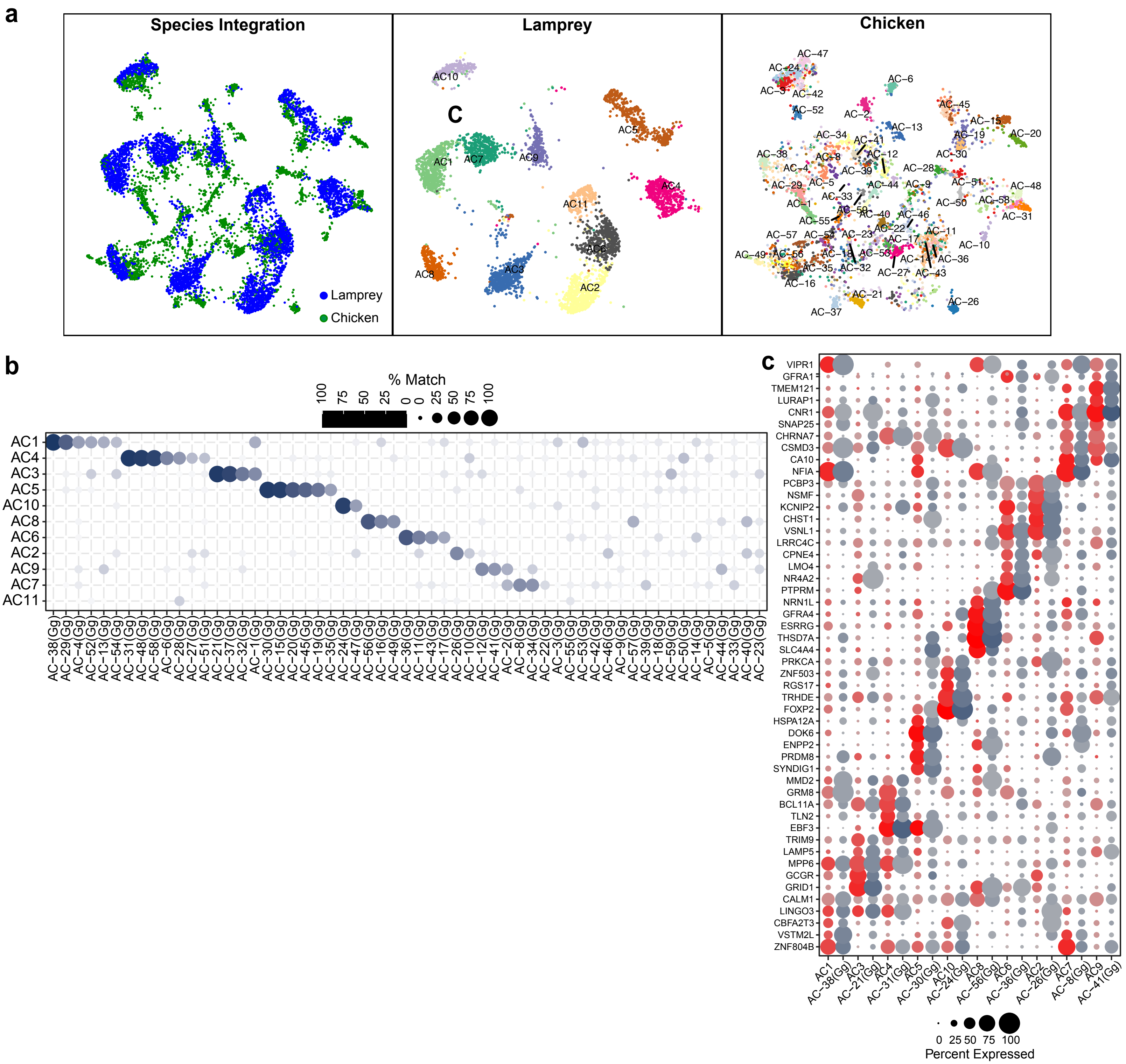

### Supplementary Figure6

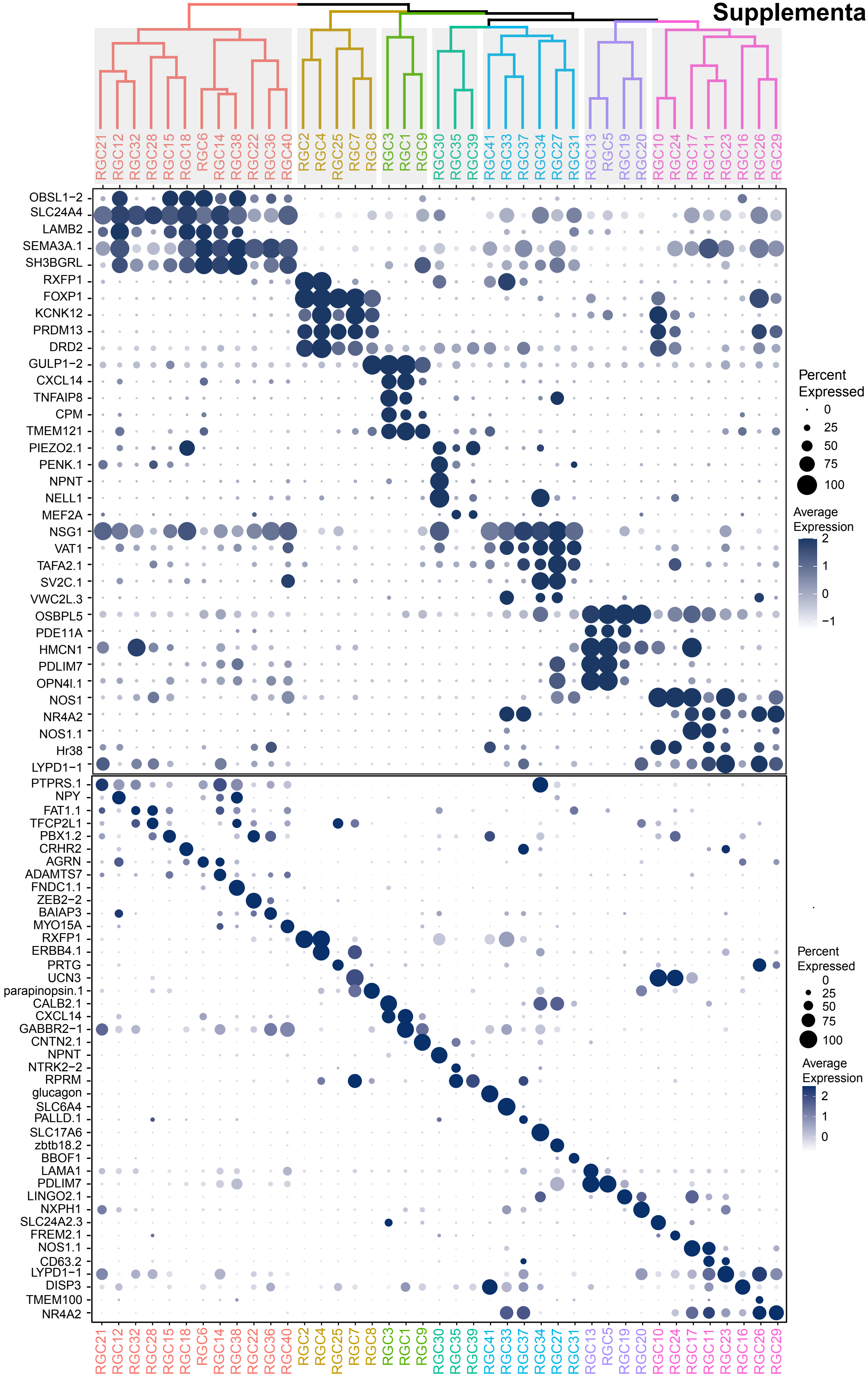
